# Parallel DNA Holliday Junctions: Myth or Reality? An Atomistic Molecular Dynamics Study

**DOI:** 10.64898/2026.07.10.737670

**Authors:** Toon Lemmens, Jiří Šponer, Petr Stadlbauer, Miroslav Krepl

**Affiliations:** Institute of Biophysics of the Czech Academy of Sciences, Královopolská 135, 612 00 Brno, Czech Republic; National Centre for Biomolecular Research, Faculty of Science, Masaryk University, Kamenice 5, 625 00 Brno, Czech Republic; Czech Advanced Technology and Research Institute, CATRIN, Palacký University, Křížkovského 511/8, Olomouc 779 00, Czech Republic

## Abstract

Holliday junctions (HJs) are key intermediates of homologous recombination and fundamental building blocks in DNA nanotechnology. Although the canonical antiparallel stacked-X conformation has been extensively characterized by X-ray crystallography, whether parallel HJ conformations exist in aqueous solution remains unresolved. Here, we address this question using extensive atomistic molecular dynamics (MD) simulations and replica-exchange umbrella sampling (REUS) free-energy calculations. Starting from canonical antiparallel HJs, standard MD simulations occasionally revealed spontaneous transitions to parallel conformations on the microsecond timescale without disrupting the DNA duplexes or passing through an open junction intermediate. REUS free-energy profiles confirmed the antiparallel state as the global minimum but also identified the parallel conformation as a well-defined local minimum, indicating it is thermodynamically metastable despite an estimated solution population below 1%. The combination of low equilibrium occupancy, microsecond interconversion dynamics, and the surprisingly close structural similarity between antiparallel and parallel junctions provides a plausible explanation for the lack of direct experimental detection. We further found that the free-energy landscape is only weakly affected by branching-point sequence and salt concentration. These results reconcile the apparent absence of experimental evidence for parallel HJs with their structural feasibility in solution and offer a fresh perspective on the historical debate.

**Graphical abstract:** 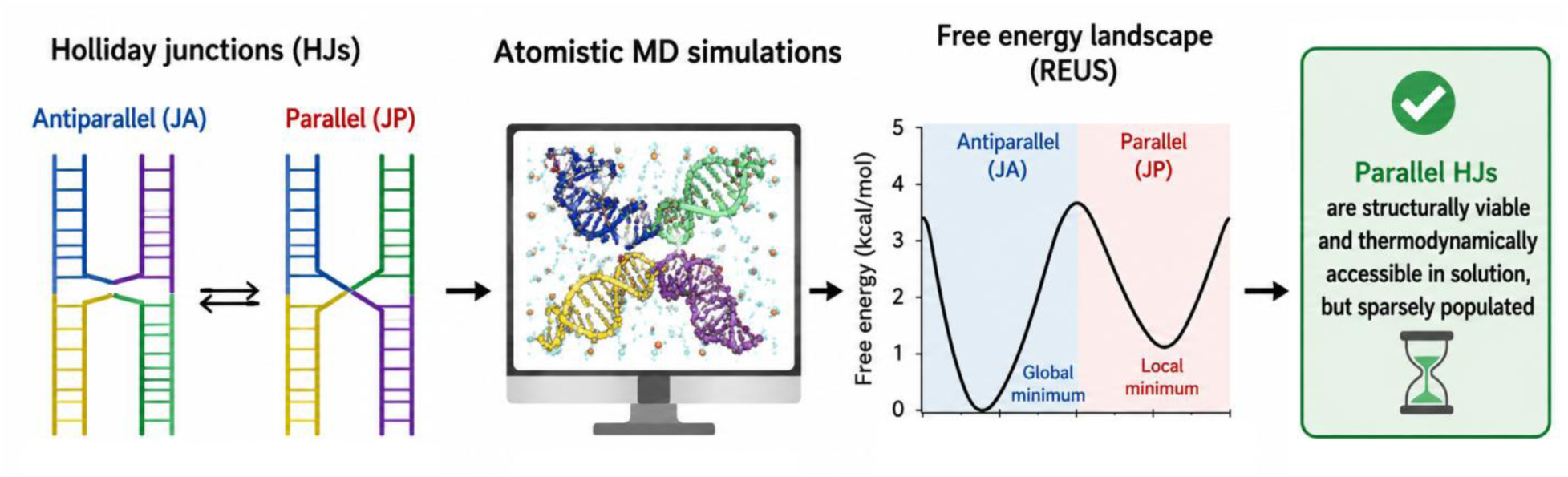

## Introduction

The Holliday junction (HJ) is a key intermediate in homologous recombination, a process in which DNA strands are exchanged between homologous DNA molecules, enabling effective repair of damaged DNA as well as promoting genetic diversity in meiotic cells (1–3). Beyond biology, the HJs form fundamental building blocks for the assembly of nanoscale architectures and machines in DNA nanotechnology (4–7). The idea of two DNA helices joined into a four-way junction was first proposed in 1964 by Robin Holliday (8). In his original depiction, the exchanging (crossover) DNA strands of the four-way junction were aligned side by side with the same 5′/3′ directionality, and the helical arms being coaxially stacked (Figure 1) (8). Holliday’s original design was later designated as the parallel HJ (JP). However, later fluorescence resonance energy transfer (FRET) experiments and X-ray structural studies rather consistently suggested that the antiparallel HJ (JA) is the dominant conformation, in which the crossover strands adopt opposite 5′/3′ directionality (Figure 1) (9–11). In fact, the JP state has not yet been directly observed in HJs by any structural biology method (11–14), prompting some studies to claim that the JP state is either not populated in solution or is exceedingly rare (15). However, one study estimated the free-energy difference between tethered HJ constructs approximately corresponding to the JA and JP states to be 1.1–1.5 kcal/mol (16). Furthermore, a recently reported process of ion-induced formation of a G-quadruplex between sequences attached to two adjacent arms of HJ suprastructures requires the HJ to first sample the JP state (17). These findings suggest that the JP state does exist in solution as a minor but accessible conformational state. However, its low population and short lifetime may place it beyond the detection limits of current structural biology methods.

**Figure 1:**
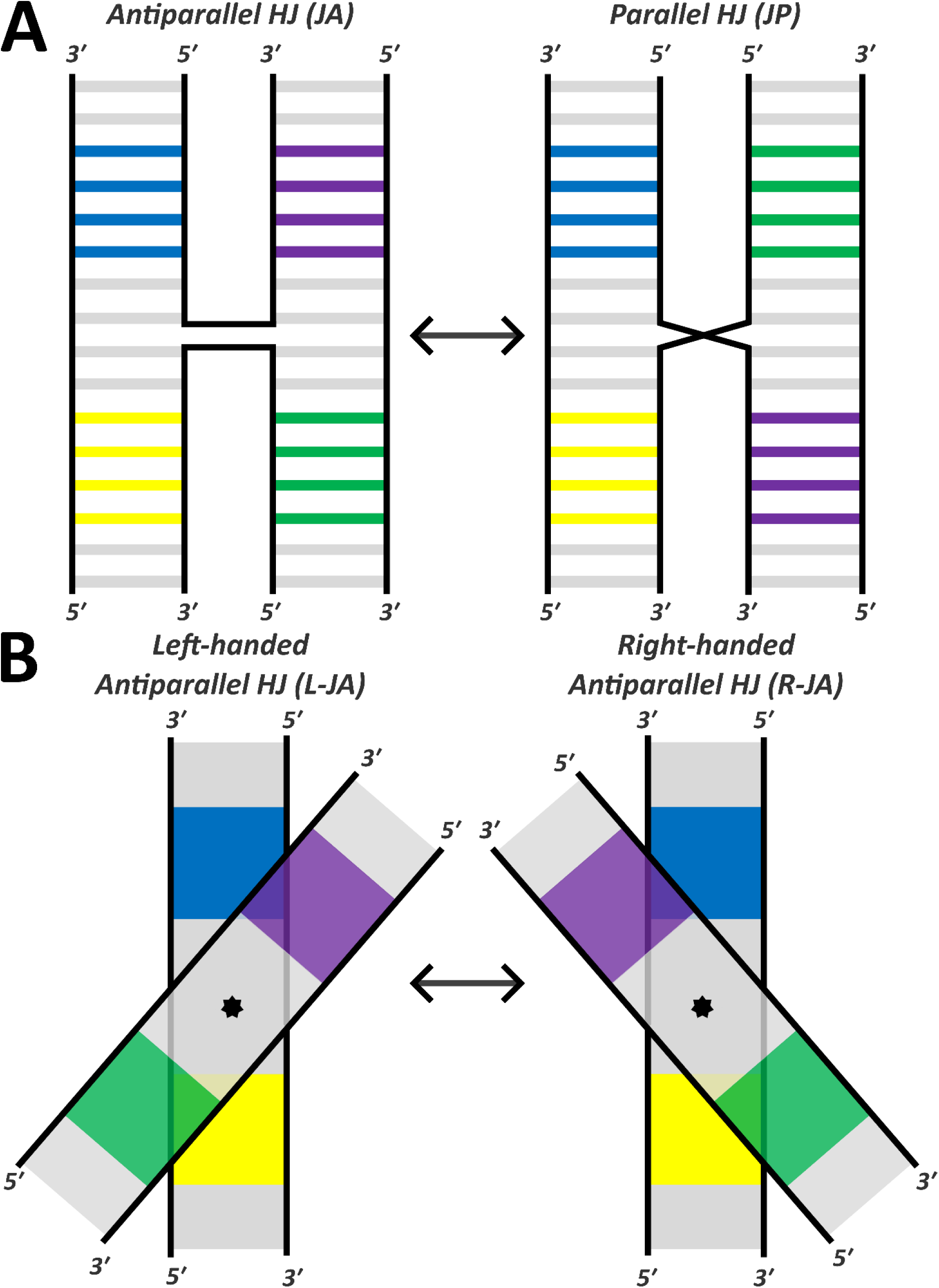
Scheme of the conformational states of the Holliday Junction. (**A**) The antiparallel (JA) and the parallel (JP) states; (**B**) A subdivision of the JA state into the left-handed antiparallel (L-JA) and the right-handed antiparallel (R-JA) state, respectively. The latter is observed in crystal structures. To better distinguish the individual states, selected base pairs in each HJ arm are color-coded.

In this study, we investigated the feasibility of the JP state of the HJ in solution using molecular dynamics (MD) simulations. MD simulations are a computational approach for probing the dynamic behavior of biomolecules at atomic resolution, based on empirically parameterized potential energy functions known as force fields (FFs). Previous studies have demonstrated the utility of MD simulations for investigating HJs (18–28). The principal advantage of the MD methodology is the effectively continuous spatial and temporal resolution while the predominant limitations stem from the FF accuracy (29–33) and the accessible simulation timescales (i.e., sampling). The latter limitation can, in some cases, be mitigated through the application of enhanced sampling methods, which aim to accelerate the relevant conformational transitions. These approaches range from “brute-force” modifications of the system’s Hamiltonian (e.g., replica exchange with solute tempering, such as REST2) (34), to knowledge-based strategies in which sampling is enhanced along a reduced set of degrees of freedom, referred to as collective variables (CVs).

Here, we performed standard MD simulations of several HJs differing in both their branching point sequence and isomeric form. When starting from the experimentally observed JA state, spontaneous transitions to the JP state can be observed in some simulations, with the JP state being relatively stable. Importantly, we observed no structural distortions of the HJ arms associated with the JP state, indicating that this state is structurally feasible. The occurrence of JA/JP transitions greatly varied in frequency and time among the individual replicates of standard MD simulations, reflecting a lack of convergence and preventing us from directly calculating the relative free energy from the unbiased MD ensembles. To obtain a rigorous estimate of the free-energy change associated with the JA/JP transition, we employed the umbrella sampling (US) method, in which the reaction coordinate undergoing the transition; in our case a dihedral angle describing the rotation of the HJ arms (the interhelical angle); is partitioned into a series of overlapping populations (windows). Each window is sampled using standard MD simulations with an applied harmonic bias potential that restrains the system to a pre-defined region of the transition. The resulting biased distributions of the CV are then combined to reconstruct the free-energy profile (potential of mean force, PMF) using reweighting methods such as the Weighted Histogram Analysis Method (WHAM) or the Multistate Bennett Acceptance Ratio (MBAR) (35–37). Due to the large number of degrees of freedom in a system of the size of the HJ, we further combined the US protocol with REST2 (referred to as REUS) (34,38–42). The PMFs obtained with REUS were highly consistent with the qualitative trends observed in unbiased standard MD simulations. Namely, the JP state, while less thermodynamically stable than JA, should be accessible as a minor state in solution. Interestingly, we observe a difference between the latest OL21 (32) and OL24 (33) DNA AMBER FFs, where the latter provides more decisive stabilization of the native JA arrangement.

Ultimately, our results suggest that while the parallel HJs as originally envisioned by Robin Holliday are not the dominant state in solution, they are accessible as a minor fluctuation state. The JA/JP interconversion occurs on the microsecond timescale, which may hinder the direct observation of the JP state in time-resolved experiments. Moreover, the JA and JP conformations are not as structurally distinct as is often assumed. In particular, their interhelical angles differ far less than would be expected for two diametrically opposed states, with the principal distinction instead being the handedness of the global supercoiling. Interestingly, the transition is smooth, diffusive, and does not require any opening of the HJ. We furthermore delineate the factors possibly contributing to the balance between the JA and JP states by showing that the branching point sequence and salt concentration do not have a significant influence. This is notable given the well-established sensitivity of the HJ structures to both factors (13,18). For example, ion concentration strongly influences the opening-closing dynamics (not explored here) of the HJs while the branching point sequence considerably influences the HJ’s ion binding properties and the ability to crystallize (18,19). Overall, our findings contribute to a more nuanced understanding of the factors governing HJ dynamics in solution by shedding light on the experimentally elusive parallel HJ states. In addition, we highlight the sensitivity of PMF calculations to the FF choice and the technical limitations of free-energy calculations.

## Materials and Methods

### Starting structures

The X-ray structure of immobile isomer I of junction J1 in the right-handed antiparallel (R-JA) state (PDB: 5KEK) was utilized as the starting structure for all MD simulations of this HJ. The structures of junction J11 in both isomers I and II were obtained by molecular modeling of the branch point base pairs using UCSF Chimera (Supplementary Figure S1) (18). For an overview of the additional characteristic conformational transitions that HJs can undergo, including opening-closing motions and isomerization, see Supplementary Figure S2. Starting structures for simulations beginning from the left-handed antiparallel (L-JA) and parallel (JP) states were manually selected from standard MD simulations in which the JA/JP transitions were observed. The selection of J1 and J11 immobile HJ sequences was motivated by those two showing highly distinct physical properties in nanotechnological application where the J1 crystallizes universally well while J11 does not form refracting crystals (18).

### System building and standard MD simulations

For all systems, the initial topology and coordinate files were generated using the tLeap module of AMBER22 (43). The DNA was described using either the OL21 (32) or OL24 FFs (33). The latter FF provides a more realistic sampling of A-DNA substates of the DNA double helices, which could be potentially relevant for the parts of the HJ immediately adjacent to the branching point. The HJs were immersed in an octahedral simulation box of SPC/E water molecules (44). In addition, the OPC (45) water model was also tested with the OL24 FF to evaluate the possible influence of the water model. A minimum distance of 13 Å between the solute and the box-boundary was used in each case. The systems were neutralized by randomly adding K^+^ ions and an excess salt concentration of 0.15 M KCl described by the Joung and Cheatham parameters was established (46). For the standard MD simulations, the established protocol (47) for nucleic acids was employed, with the pmemd.MPI program used for equilibration. The production simulations were then calculated on GPUs using the pmemd.cuda (48), and three replicates of each system were simulated for 10 μs at 300 K. An integration step of 4 fs was enabled using SHAKE (49) and hydrogen mass repartitioning (50). Long range electrostatics were handled by the periodic boundary condition using the particle mesh Ewald method (51). The cut-off distance for the Lennard-Jones (LJ) interactions was 9 Å, and temperature and pressure were regulated using the Langevin thermostat (52) and Monte Carlo barostat (39), respectively. To prevent base pair fraying, a 2 kcal/mol stabilizing structure-specific HBfix (53) was applied between acceptor atoms and hydrogens of the H-bonds constituting the terminal base pairs of each HJ arm (four base pairs in total).

### Definition of the collective variable (CV) for describing the JA/JP transition of the HJ

The CV was defined as a dihedral angle whose four defining points were given by centers of mass (COM) located at specific positions along each HJ arm, equidistant from the junction’s branching point (Figure 2A). Rotation along this dihedral (henceforth referred to as the CV) closely captures the JA/JP transition process. A highly non-trivial technical complication arises from the CV values corresponding to the R-JA and JP states partially overlapping within the periodic space of the dihedral (Figure 2B). This occurs due to the HJ (as with any DNA helix) supercoiling when twisted too far in any particular direction. Unfortunately, the free-energy minima of both the R-JA and JP states – the respective end-points of the JA/JP transition – lie very close to the negative and positive supercoiling regions, respectively. Hence, restricting the range of the CV to avoid the supercoiling regions is not feasible as it would severely bias the results. We also note that despite significant effort, we were unable to identify a secondary or alternative CV that would *unambiguously*, at all times distinguish between the negative and positive supercoiling states. Unfortunately, this precluded the use of free-energy methods that reconstruct the PMF from trajectory frames without accounting for their history (e.g., metadynamics (54) or OPES (55)), as identical biasing potentials would inevitably be assigned to structurally highly distinct yet according to the CV technically overlapping regions. Thus, instead, we opted to apply umbrella sampling (US) combined with REST2 exchanges between the umbrella windows (REUS method) (Figure 2) (34,38–41). In US calculations, each sampling window is confined to a relatively narrow range of CV values, so that the overlapping supercoiling regions can be reliably and easily distinguished during the reweighting process.

**Figure 2:**
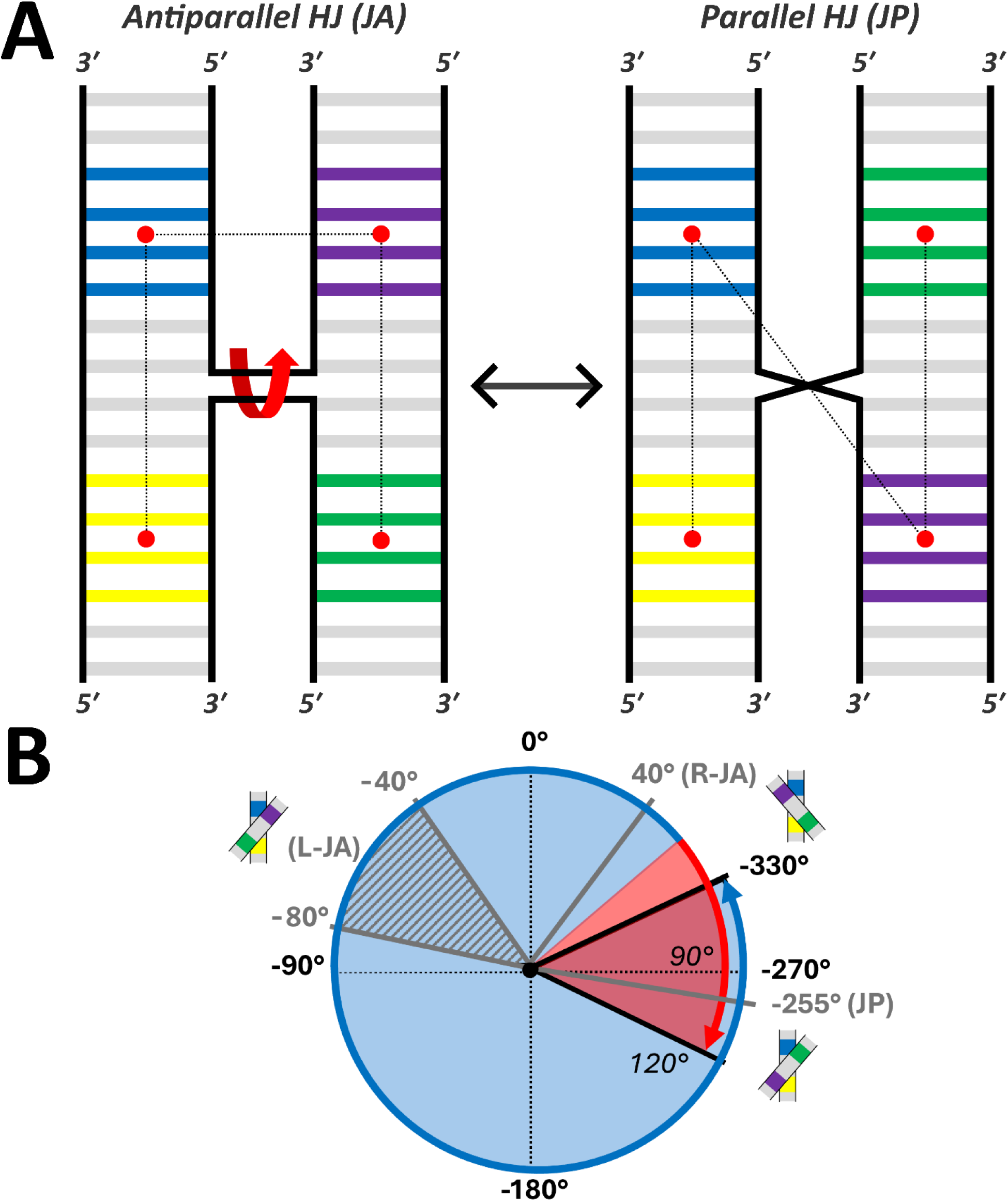
The CV describing the JA/JP transition of HJ. (**A**) The dihedral angle defining the CV is indicated by the dotted line connecting the four red points. Each point is defined as a center of mass (COM) of C1′ atoms of four base pairs located equidistantly from the branching point within each HJ arm. The residues included in the calculation of the COMs are highlighted in distinct colors for each arm, with the same color-coding used consistently throughout the study. (**B**) A scheme illustrating the position of individual HJ states (i.e., R-JA, L-JA and JP) within the one-dimensional space of the CV. The approximate location of the free-energy minima corresponding to each state are indicated with grey lines. The location of the L-JA is better described as an interval, indicated by the hatched area with grey lines. It is essential to note that due to structural supercoiling, the rotation cannot proceed infinitely in one direction. The solid black lines indicate the approximate supercoiling limits beyond which the free energy increases sharply. Lastly, note that the fluctuation ranges of the end point states (R-JA and JP) partially overlap within the periodic space, requiring the use of unwrapped dihedral function to differentiate them when reconstructing the PMFs. This is indicated by the partially overlapping red and blue parts of the circle.

### Umbrella sampling simulations with REST2 (REUS)

The REUS calculations were initiated from the R-JA structure of the HJ, possessing a CV value of ∼50°. Each window of the calculations was sampled for 100 ns and spanned 10° confined by harmonic restraint of 30 kcal×mol^-1^×rad^-2^. The windows were propagated over the range of -330° to 120° as depicted in Figure 2B (47 windows in total, corresponding to a cumulative sampling time of 4.7 µs per single transition). The equilibration and production simulation setup was consistent with that used for standard MD simulations (see above), except that the REUS simulations were carried out under canonical ensemble conditions. At least three independent replicates were performed for each setup. The REST2 protocol was implemented using 12 replicas, with the scaling factor λ ranging from 1 to 0.8344, yielding an exchange acceptance rate of 25% on average. The entire HJ molecule (64 nucleotides) was included in the solute scaling (i.e., defined as the “hot” region), and exchange attempts between replicas were performed every 10 ps. The CV was not scaled across replicas. The solute-tempering replica-exchange protocol was employed primarily to smooth out various stochastic inter-arm interactions that could randomly arise during the transition rather than to accelerate the transition itself. Note that the JA/JP transition is somewhat unusual in terms of its free-energy profile. Namely, it is best described as a slow diffusion along an extended coordinate rather than as a process governed by sharply defined free-energy barriers. In other words, its greatest slowdown factor is the stochastic, back-and-forth nature of the diffusion rather than the need to overcome a sizable free-energy barrier. This makes the REUS an optimal method to study this transition as the random walk nature of the diffusion is intrinsically avoided by the US protocol.

To prevent excessive bending of the HJ arms, which could potentially interfere with the JA/JP transition, we introduced two auxiliary angular restraints for the REUS calculations. Each angle was defined using the two centers of mass (COMs) along a continuous HJ helix that are also being used to define the CV (Figure 2A; i.e. yellow to blue and purple to green), with the middle point defined by the COM of the C1′ atoms of the eight nucleotides closest to the branch point (Supplementary Figure S3). A lower-wall boundary of 135° was imposed, with smaller angles linearly penalized using a restraint with a force constant of 300 kcal×mol^-1^×rad^-2^. These restraints were introduced to prevent the helices from bending excessively under the applied CV bias and thereby escaping the JA/JP transition pathway through an orthogonal mode of motion. We emphasize that angles below 135° were rarely sampled in standard unbiased MD simulations during the spontaneous JA/JP transitions (Supplementary Figure S4) and we thus suggest these angular restraints do not significantly affect the PMFs.

### Analysis

In all trajectories (standard MD and REUS), we monitored the development of the CV (see above) using cpptraj. All standard MD trajectories were visually inspected using VMD, and the stability of base pairs was monitored visually and with cpptraj. In the REUS calculations we monitored all the replicas visually and with cpptraj while the lowest (unscaled) replica was used to derive the free-energy profiles (PMFs). The sampled CV values were saved every 10 integration steps for each of the windows. The existence and extent of the overlap between the adjacent windows (an essential condition of obtaining physically meaningful PMFs with REUS calculations) were extensively monitored by analyzing the histograms of the sampled CV values per window (Supplementary Figures S5-S12). The PMFs were calculated using either the WHAM (35) or MBAR (37) methods. The principal difference between the reweighting methods is that MBAR does not require binning of the sampled energies and therefore avoids the discretization bias associated with histogram-based approaches such as WHAM (37). Consequently, MBAR typically makes more efficient use of the available data and can provide more precise free-energy estimates. In practice, however, the PMFs obtained using the two methods were nearly identical, with differences typically smaller than the estimated statistical uncertainty. Individual PMFs were calculated for each replicate and subsequently averaged to obtain the final PMF. All plots were generated using python’s matplotlib module. For clarity, we do not display the first and last 10° in each PMF graph as the free energy rises sharply beyond those points. To make the graphs comparable, we also cap the displayed PMFs at +10 kcal/mol.

## Results and Discussion

In this work, we present results of standard and enhanced sampling MD simulations examining the structure and dynamics of three immobile HJs undergoing the JA/JP transition. In immobile junctions, the branching point sequence prevents branch migration (Supplementary Figure S2) by design (18). Using an extensive set of standard (unbiased) MD simulations, we show that when initiated from the native JA state, the JP state is accessible on the 10-microsecond timescale. Furthermore, the JP state is not associated with any structural distortions of the helical arms, indicating that the JP state is fully feasible from a structural standpoint, i.e., it is compatible with the HJ structure. We further characterize the JA/JP transition using enhanced-sampling free-energy REUS calculations (see Methods), which capture the one-dimensional free-energy profile (PMF) associated with the transition. This computational framework was applied to two immobile HJs, J1 and J11, differing in their branching-point sequences (Supplementary Figure S1). In addition, in the case of J11, both isomer I and II were examined. To describe the DNA, we applied the two latest AMBER DNA FF versions, OL21 (32) and OL24 (33). OL24 is recommended by the current AMBER reference manual (56) while the OL21 was the preceding recommended version (57). We also assessed the effects of the water model and salt concentration. The total simulation time presented in this work amounts to ∼667 μs (Table 1). For all HJs, the endpoints of the JA/JP transition are the right-handed antiparallel (R-JA) and right-handed parallel (JP) states, characterized by CV values of ∼40° and ∼−255°, respectively. An intermediate left-handed antiparallel state (L-JA) is also located along the transition pathway (Figure 1). While formally just an intermediate, the L-JA state does have a sufficient population in the simulations to warrant its consideration as a separate state in the analyses. Its corresponding CV values range from ∼−40° to ∼−80°. In contrast, no stable left-handed parallel state (L-JP) was observed, and only the right-handed parallel conformation was sampled. We therefore refer to the parallel state simply as JP.

**Table 1:** List of the standard MD and REUS simulations.^a^.

| HJ / DNA FF | Replicas × Length ($\mu\text{s}$) |
| --- | --- |
| **Standard MD – started from R-JA** |  |
| J1/OL21 | $3 \times 10$ |
| J11-IsoI/OL21 | $3 \times 10$ |
| J11-IsoII/OL21 | $3 \times 10$ |
| J1/OL24 | $3 \times 10$ |
| J11-IsoI/OL24 | $3 \times 10$ |
| J11-IsoII/OL24 | $3 \times 10$ |
| **Standard MD – started from L-JA** |  |
| J1/OL21 | $3 \times 10$ |
| J11-IsoI/OL21 | $3 \times 10$ |
| J11-IsoII/OL21 | $3 \times 10$ |
| J1/OL24 | $3 \times 10$ |
| J11-IsoI/OL24 | $3 \times 10$ |
| J11-IsoII/OL24 | $3 \times 10$ |
| **Standard MD – started from JP** |  |
| J1/OL21 | $3 \times 10$ |
| J11-IsoI/OL21 | $3 \times 10$ |
| J11-IsoII/OL21 | $3 \times 10$ |
| J1/OL24 | $3 \times 10$ |
| J11-IsoI/OL24 | $3 \times 10$ |
| J11-IsoII/OL24 | $3 \times 10$ |
| **REUS<sup>b</sup>** |  |
| J1/OL21 | $4 \times 4.7$ |
| J1/OL21-1M-KCl<sup>c</sup> | $3 \times 4.7$ |
| J11-IsoI/OL21 | $4 \times 4.7$ |
| J11-IsoII/OL21 | $4 \times 4.7$ |
| J1/OL24 | $3 \times 4.7$ |
| J1/OL24-OPC<sup>d</sup> | $3 \times 4.7$ |
| J11-IsoI/OL24 | $3 \times 4.7$ |
| J11-IsoII/OL24 | $3 \times 4.7$ |

<sup>a</sup>The name of the HJ structure reflects the difference in branching-point base-pair sequence (J1 and J11, see Supplementary Figure S1) and the corresponding isomer (isomer I or II – relevant only for J11). Only isomer I was studied for J1 and is therefore not specified in its name; see Methods. R-JA, L-JA and JP refer to the right-handed antiparallel, left-handed antiparallel and parallel starting structures being used. See Methods for further explanation.
<sup>b</sup>Each REUS calculation consisted of 47 windows of 100 ns each, corresponding to a cumulative simulation time of 4.7 μs per calculation.
<sup>c</sup>Simulation was conducted with excess salt concentration of 1 M KCl instead of 0.15 M.
<sup>d</sup>OPC water model was used instead of SPC/E.

### Standard MD simulations of the J1 detail the JA/JP transition process

In our initial standard MD simulations of the J1 structure using the OL21 FF, one replicate exhibited a complete irreversible transition from the starting R-JA state to the JP state. In the remaining two replicates, reversible fluctuations between the R-JA and L-JA states were observed (Figure 3, top).

**Figure 3:**
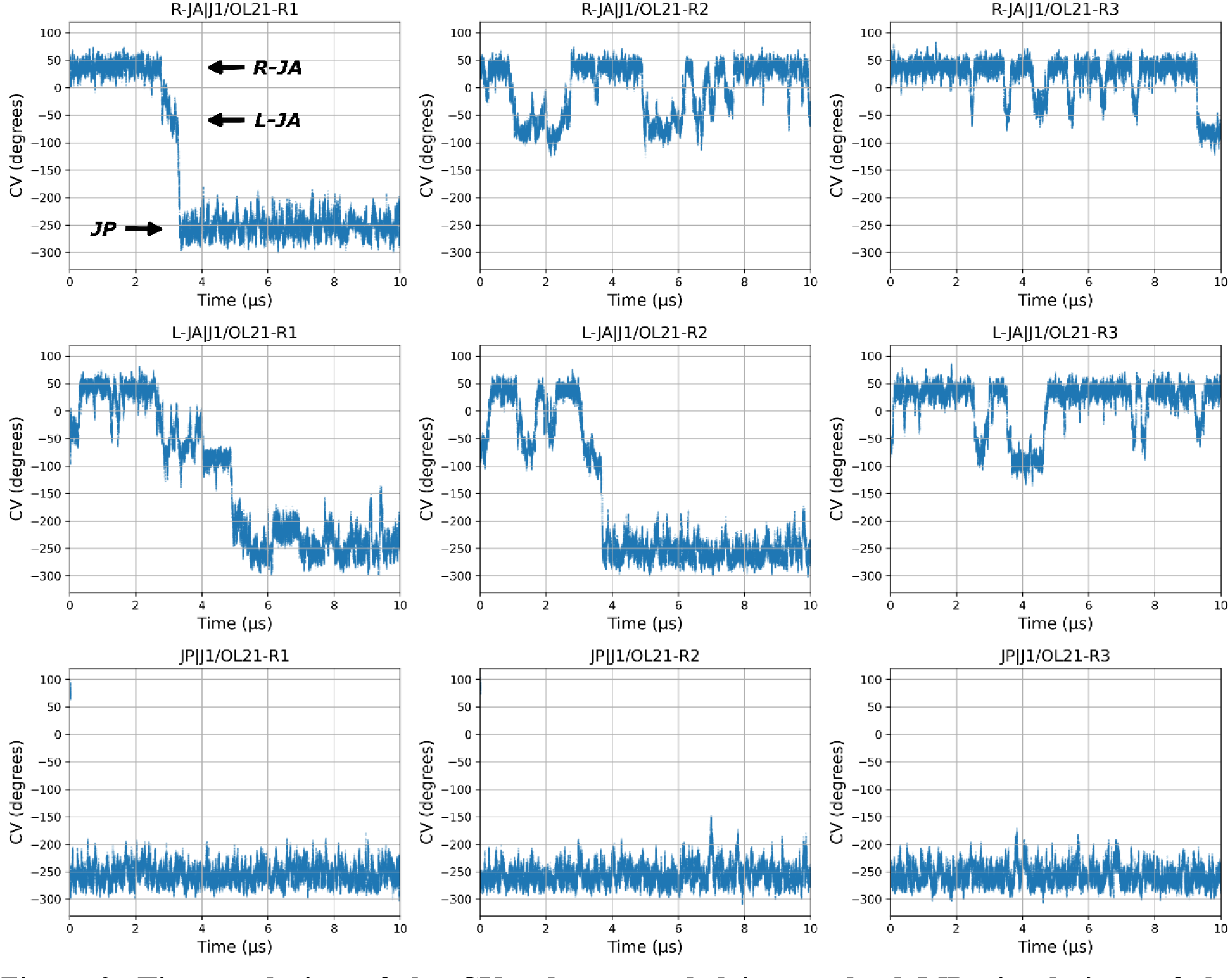
Time evolution of the CV values sampled in standard MD simulations of the J1/OL21 system. Top, middle and bottom rows show the three standard MD simulation replicates initialized from the R-JA, L-JA and JP states, respectively. The arrows in the upper left graph mark the CV values corresponding to the R-JA, L-JA and JP states respectively. For the average CV values of each state, see Supplementary Table S1. Equivalent analysis for both isomers of J11 is presented in the Supplementary Figures S13 and S14.

Throughout these transitions, the open HJ state is not sampled. Based on these simulations, representative L-JA and JP conformations were selected as starting structures for additional simulations. In the subsequent simulations performed with the OL21 FF, the JP starting state remained universally stable, whereas the intermediate L-JA starting state eventually transitioned either to the R-JA state, with reversible excursions back to L-JA, or to the JP state (Figure 3). With OL24, the balance of the states seems to be shifted more in favor of the native R-JA arrangement relatively to OL21. Specifically, no transitions to the JP state were observed on the simulation timescale when starting from the R-JA state, while transitions to the L-JA state were also substantially less frequent compared to the OL21 simulations. Simulations initiated from the L-JA state exhibited rapid relaxation to the R-JA state. In contrast, simulations initiated from the JP state showed a single transition back to the R-JA state in one replicate, whereas the JP state remained stable in the other two replicates (Figure 4). Within the limits of the present sampling, these observations suggest that the OL24 FF stabilizes the R-JA state more strongly than OL21. Importantly, both tested FFs qualitatively support kinetically stable antiparallel and parallel states with the lifetimes on a scale of ∼10 µs. However, the number of observed transitions is entirely insufficient to draw any further quantitative conclusions from the standard MD simulations alone. At most, only a single irreversible JA/JP transition was observed within the timescale of an individual simulation, highlighting substantial sampling limitations. However, an important qualitative finding is that the JP conformation does not seem to be destabilized by any visible strain.

**Figure 4:**
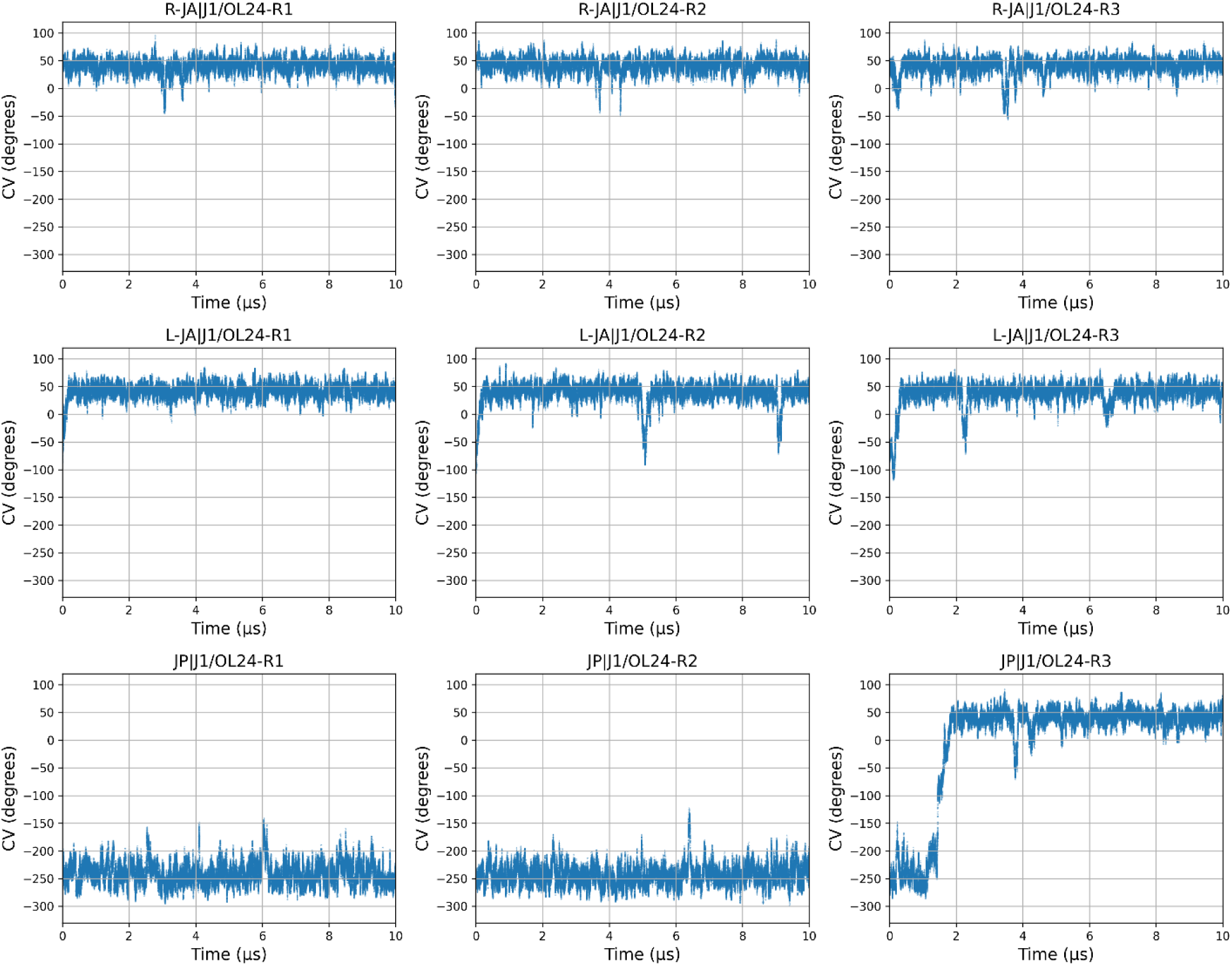
Time evolution of the CV values sampled in standard MD simulations of the J1/OL24 system. Top, middle and bottom rows show the three standard MD simulation replicates initialized from the R-JA, L-JA and JP states, respectively. For the average CV values of each state, see Supplementary Table S1. Equivalent analysis for both isomers of J11 is presented in the Supplementary Figures S15 and S16.

### J11 simulations reveal only minor differences in the JA/JP transition

Immobile junction J11 differs from J1 solely in the sequence of the base pairs adjacent to the branching point (Supplementary Figure S1). Standard MD simulations of J11 revealed only minor differences in the JA/JP transition process described above (Supplementary Figures S13 to S16), at least at a qualitative level and particularly in light of the clear sampling limitations. The only notable difference that appears robust is the relatively higher population of the intermediate L-JA state in J11 isomer I compared to either J11 isomer II or J1. We tentatively suggest this increased anisotropy within the global JA-state region might be one of the factors contributing to poor crystallization of the J11 isomer I (18). Overall, these observations suggest that neither the HJ branching-point sequence nor the specific isomer substantially affects the JA/JP transition itself, although both factors may influence the local dynamics within the individual R-JA, L-JA, and JP states. The difference between OL24 and OL21 FFs was also, within the limits of sampling, the same as observed for J1 (see above and Figure 3 and 4). This includes the greater overall stability of the R-JA state with OL24 and somewhat reduced interconversion dynamics between the R-JA and L-JA states (Supplementary Figure S15 and S16).

### PMFs of the JA/JP transition characterized using REUS calculations

Although the standard MD simulations provided detailed insight into the transition process (see above), the individual trajectories did not reveal a sufficient number of transitions to reach thermodynamically converged R-JA, L-JA, and JP populations. This is not unexpected, as the JA/JP transition appears to be quite diffusive in nature, with the HJ arms having to chiefly overcome friction arising from the explicit solvent environment, with back-and-forth attempts and unsuccessful transitions aborted before completion. As a result, transitions are relatively sluggish and infrequent. To obtain quantitative information on the transition process, we performed REUS calculations using the CV to describe the JA/JP transition. The design of the CV and the choice of enhanced sampling methodology resulted from an extensive search for an optimal simulation protocol. Initially, we tried to perform well-tempered metadynamics calculations, however, this approach turned out to be inapplicable due to the periodicity problem connected to the CV where the end-states partially overlap (see Methods and Figure 2B), leaving umbrella sampling as the most straightforward and appropriate methodology to tackle the issue. The resulting free-energy profiles were calculated using either the MBAR or WHAM reweighting approaches. We present both estimators below; however, the resulting free-energy profiles are highly similar, with the observed differences falling within the known limitations and uncertainty of enhanced-sampling methodologies (Figures 5 and 6) (19,58).

**Figure 5:**
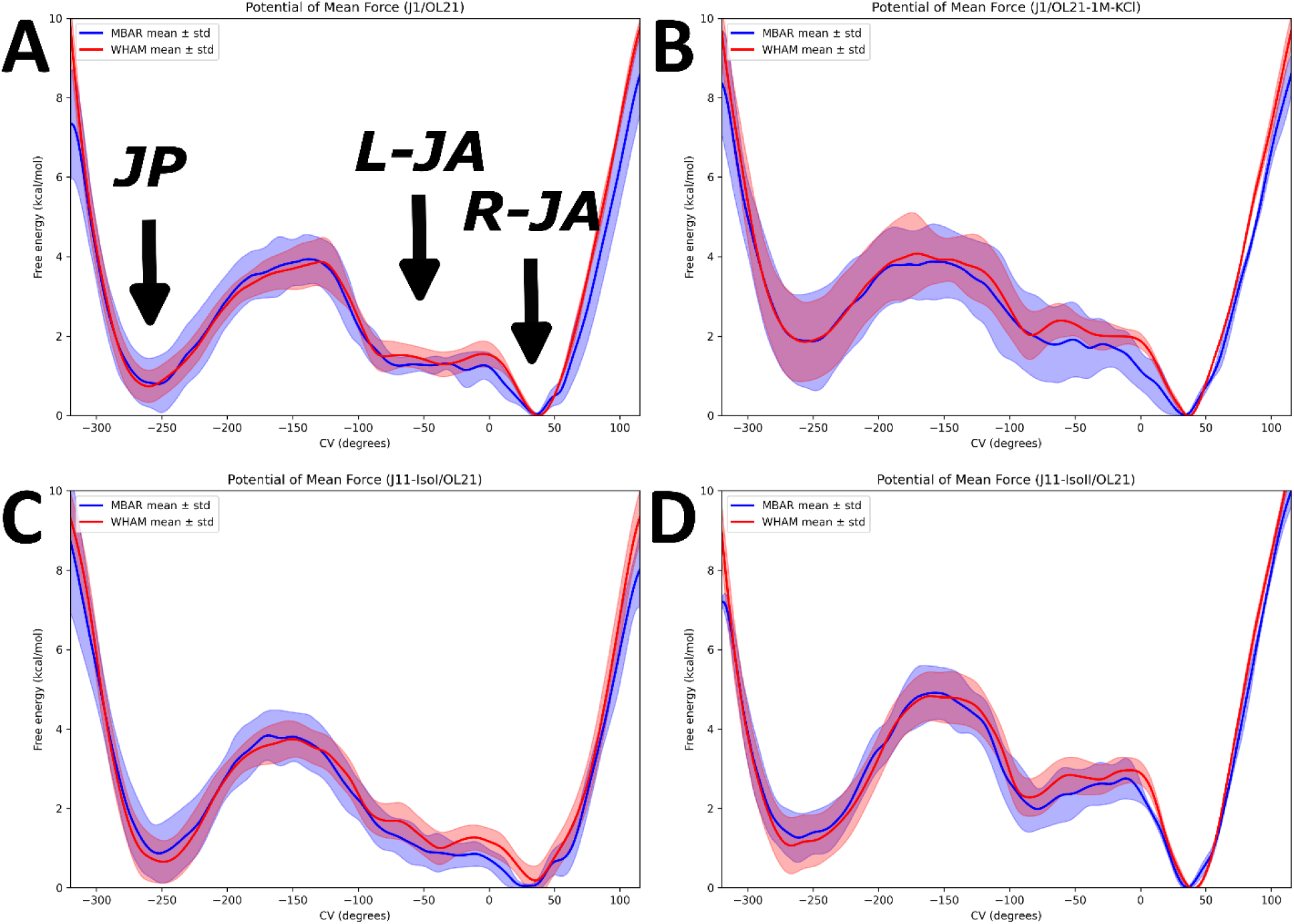
PMFs from REUS calculations using OL21 FF. Profiles averaged from three replicates of the REUS calculations obtained using the MBAR and WHAM reweighting estimators are presented in blue and red, respectively. Their standard deviation across the replicates is shown as a transparent cloud. In the first graph, we indicate the locations of the free-energy minima corresponding to the R-JA, L-JA and JP states with black arrows. (A) J1/OL21, (B) J1/OL21 with 1 M KCl concentration, (C) J11-isoI/OL21, and (D) J11-isoII/OL21.

**Figure 6:**
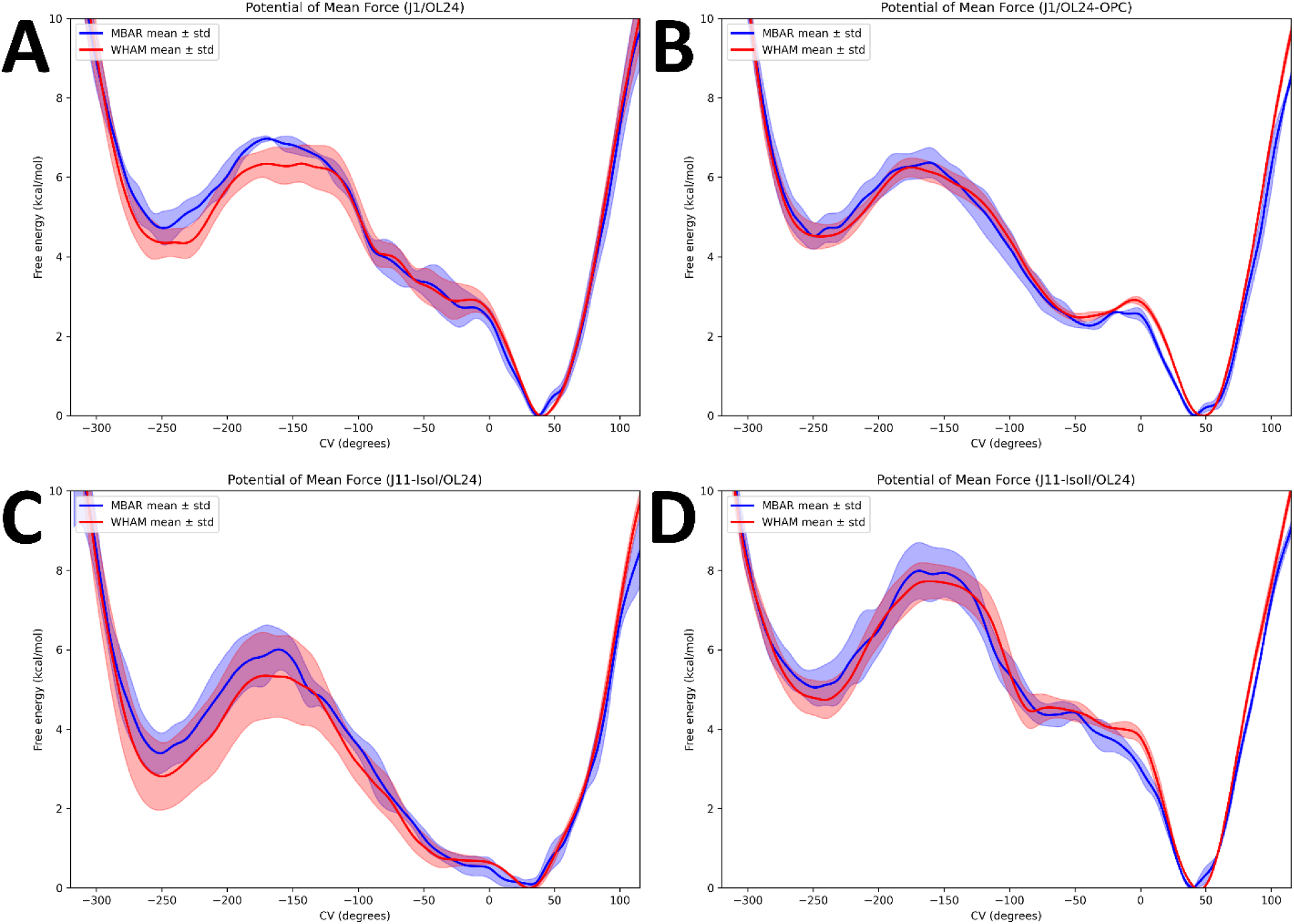
PMFs from REUS calculations using OL24 FF. Profiles averaged from three replicates of the REUS calculations obtained using the MBAR and WHAM reweighting estimators are presented in blue and red, respectively. Their standard deviation across the replicates is shown as a transparent cloud. (A) J1/OL24, (B) J1/OL24 combined with OPC water model, (C) J11-isoI/OL24, and (D) J11-isoII/OL24.

### REUS identifies the R-JA state as the global free-energy minimum for both J1 and J11 using the OL21 FF

The free-energy profiles (PMFs) of the JA/JP transition for J1 exhibit two distinct minima at CV values of approximately 40° and −255°, corresponding to the R-JA and JP states, respectively (Figures 2 and 5). These are fully consistent with the values observed in standard MD simulations (Figures 3 and Supplementary Table S1). The free-energy difference between the R-JA and JP states is estimated to be ∼1 kcal/mol (Table 2), making the JP state only modestly less stable within the limits of the sampling. Their corresponding two minima are separated by a broad free-energy barrier of ∼4 kcal/mol (Table 2) which tops around CV values of −150°. In contrast, the L-JA state forms only a shallow minimum that is more appropriately described as a broad free-energy plateau, with CV values spanning approximately from −80° to 0° (Figure 5). This is once again consistent with observations from the standard MD simulations which show extensive fluctuations within this region (Figure 3). Highly similar free-energy profiles were obtained also for the J11 in both isomers (Figure 5C and 5D), suggesting that the balance between the R-JA and JP states is relatively insensitive to the identity of the branching-point base pairs. Notably, the R-JA state remains the global minimum for all simulated junctions, indicating that the experimentally observed difficulty in crystallizing the isomer I of J11 (and some other sequences not tested here) cannot be rationalized by a potentially increased population of the JP state (18).

**Table 2:**
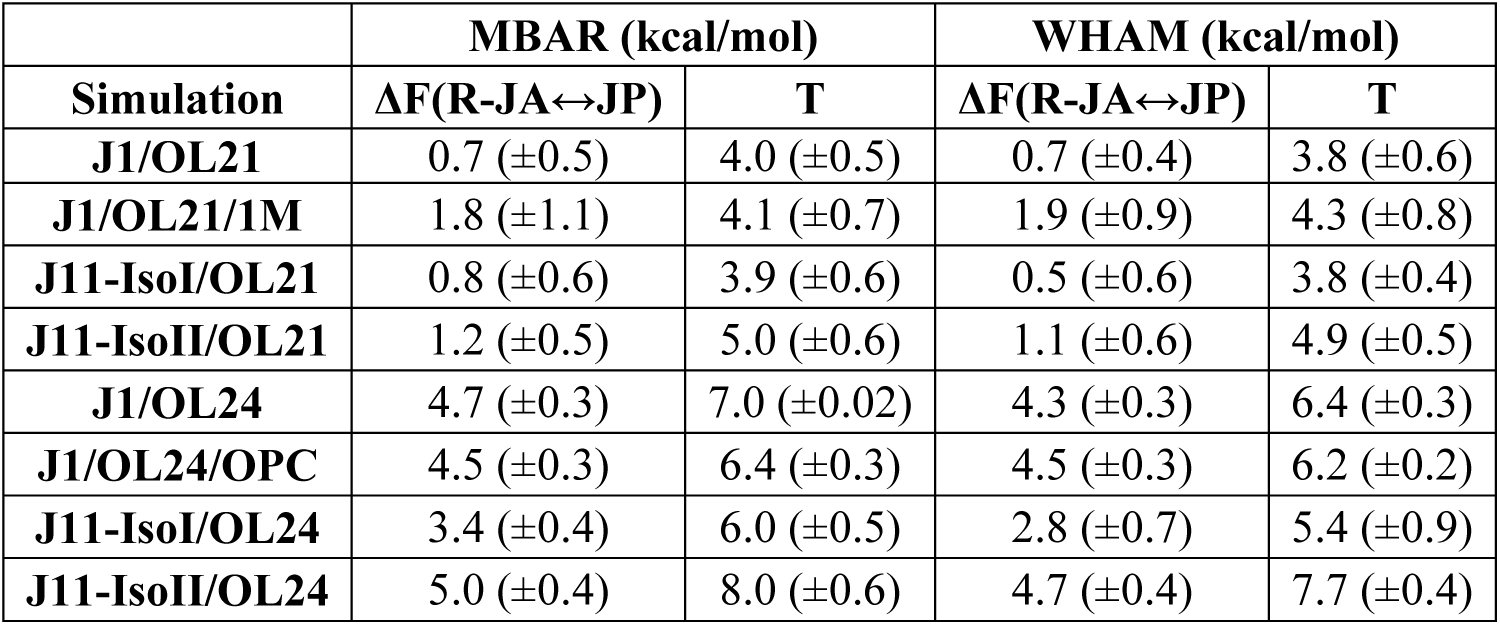
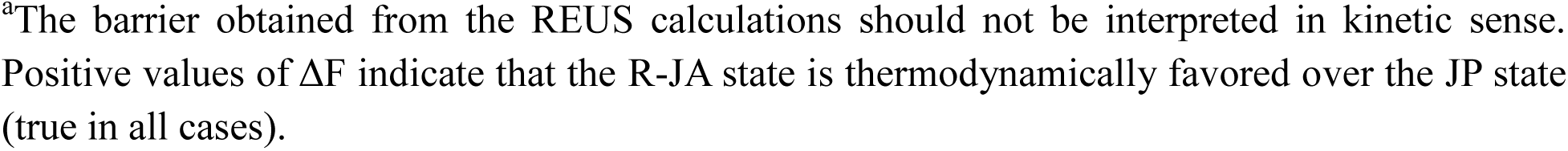
The free-energy differences between R–JA and JP states (ΔF), and the height of barrier (T) in REUS calculations.^a^.

| Simulation | MBAR (kcal/mol) |  | WHAM (kcal/mol) |  |
| --- | --- | --- | --- | --- |
| | $\Delta F(\text{R-JA} \leftrightarrow \text{JP})$ | T | $\Delta F(\text{R-JA} \leftrightarrow \text{JP})$ | T |
| J1/OL21 | 0.7 ($\pm 0.5$) | 4.0 ($\pm 0.5$) | 0.7 ($\pm 0.4$) | 3.8 ($\pm 0.6$) |
| J1/OL21/1M | 1.8 ($\pm 1.1$) | 4.1 ($\pm 0.7$) | 1.9 ($\pm 0.9$) | 4.3 ($\pm 0.8$) |
| J11-IsoI/OL21 | 0.8 ($\pm 0.6$) | 3.9 ($\pm 0.6$) | 0.5 ($\pm 0.6$) | 3.8 ($\pm 0.4$) |
| J11-IsoII/OL21 | 1.2 ($\pm 0.5$) | 5.0 ($\pm 0.6$) | 1.1 ($\pm 0.6$) | 4.9 ($\pm 0.5$) |
| J1/OL24 | 4.7 ($\pm 0.3$) | 7.0 ($\pm 0.02$) | 4.3 ($\pm 0.3$) | 6.4 ($\pm 0.3$) |
| J1/OL24/OPC | 4.5 ($\pm 0.3$) | 6.4 ($\pm 0.3$) | 4.5 ($\pm 0.3$) | 6.2 ($\pm 0.2$) |
| J11-IsoI/OL24 | 3.4 ($\pm 0.4$) | 6.0 ($\pm 0.5$) | 2.8 ($\pm 0.7$) | 5.4 ($\pm 0.9$) |
| J11-IsoII/OL24 | 5.0 ($\pm 0.4$) | 8.0 ($\pm 0.6$) | 4.7 ($\pm 0.4$) | 7.7 ($\pm 0.4$) |

<sup>a</sup>The barrier obtained from the REUS calculations should not be interpreted in kinetic sense. Positive values of $\Delta F$ indicate that the R-JA state is thermodynamically favored over the JP state (true in all cases).

### The OL24 FF further stabilizes the global free-energy minimum corresponding to the R-JA state

In contrast to the REUS simulations performed with the OL21 FF, in which the R-JA state was more stable than the JP state by only ∼1–2 kcal/mol, the OL24 FF predicts a substantially larger free-energy difference of ∼4.5 kcal/mol in favor of the R-JA state, with the free-energy barrier reaching ∼7 kcal/mol at its maximum (Figure 6, Table 2). In addition, the L-JA state also appears less stable in OL24 compared to OL21. These results are fully consistent with the greater stability of the R-JA state observed in the standard MD simulations and R-JA/L-JA transitions appearing more rarely (Figure 4 and 6). Nevertheless, the JP state still forms a clearly defined local free-energy minimum on the PMF (Figure 6), and the JP state can remain stable in standard MD on the 10-µs time scale (Figure 4 and Supplementary Figures S15 and S16). The differences between the J1 and J11 junctions are relatively minor and follow the same trends with the OL24 FF as with OL21 (Figures 5 and 6), most notably the greater accessibility of the L-JA state in J11-isoI (Figure 6C).

### Salt conditions and water-model selection do not significantly affect the JA/JP transitions

Cation concentration strongly influences biophysical properties of DNA HJs, most notably their opening and closing dynamics (not explored in the present work, see also Supplementary Figure S2) (19). We therefore investigated whether a substantially increased salt concentration (1 M of KCl) could also affect the JA/JP transition in some fashion, even though the junction remains entirely closed during the process. The REUS simulations performed under such high-salt conditions yielded free-energy profiles similar to those obtained under the standard 0.15 M KCl conditions, with the JP state being only slightly destabilized (Figure 5B and Table 2). These results suggest that ion concentration has only a limited effect on the JA/JP transition. We avoided testing Mg^2+^ ions because of their well-known FF and sampling limitations. Nevertheless, we suggest that they would have a similarly limited effect on the JA/JP transition. Indeed, salt concentration appears to primarily influence the opening-closing dynamics rather than the JA/JP transition, which proceeds entirely through the closed state without junction opening. Accordingly, sufficiently low ion concentrations would be expected to impede the JA/JP transition by shifting the equilibrium toward the open state.

Secondly, given the predominantly diffusive nature of the JA/JP transition, for which solvent friction is expected to constitute the primary obstacle, we hypothesized that the choice of explicit-solvent water model could influence the free-energy landscape. To verify this, we performed REUS calculations using the OL24 FF in combination with the OPC water model. However, this setup again produced only minor differences compared to calculations performed with the SPC/E water model. In particular, both the estimated free-energy difference between the R-JA and JP states and the overall shape of the PMF remained highly similar (Figure 6B and Table 2). These results suggest that the calculated PMFs associated with the JA/JP transition are not significantly affected by the choice of water models.

### JA and JP states *are not* related by two-fold symmetry

In the literature, the JA/JP transition of the HJ is traditionally conceptualized as a 180° rotation of one duplex relative to the other. For instance, the JA and JP states can be schematically described with idealized interhelical angles of 0° and 180°, respectively (Figure 7A). For illustrative purposes, we also adopted this simplified representation in Figure 1A. However, our calculations clearly reveal that the actual JA/JP transition differs substantially from this simplified model. Most notably, the free-energy minima corresponding to the dominant antiparallel R-JA and the JP states are separated by ∼300° along the interhelical angle coordinate, corresponding to an almost full circle rotation rather than a simple 180° inversion of a duplex (Figure 7B). Consequently, the R-JA and JP states become formally close when the handedness of the structural supercoiling is ignored. In fact, when projected on the interhelical angle (used as CV in our calculations), there is a non-negligible fluctuation overlap between the two states observed in standard MD simulations (Supplementary Figure S17). This surprising feature of the JA/JP transition posed a major challenge during the design of the CV used for the free-energy calculations (see above and the Methods). We suspect that it could also complicate the interpretation of classical FRET experiments often performed with HJs (10,13,15,20,59–61) in which fluorophores are attached to HJ arms. In fact, our results show the R-JA and JP states primarily differ in supercoiling and can partially overlap in other low-dimensionality distance and angular descriptions. The combination of the ∼10-microsecond timescale lifetimes, flexibility of the attached fluorophores, and the relatively low expected population of the JP state could potentially obscure the difference between the R-JA and JP states in time-resolved experiments. Lastly, we note an interesting asymmetry of the JA/JP transition where the distinct left-handed free-energy minimum exists exclusively within the antiparallel region (the L-JA state) whereas only the right-handed conformation forms a well-defined free-energy minimum in the JP region. In fact, the hypothetical L-JP state would be expected at CV values coinciding with the highest point of the free-energy barrier along the JA/JP transition pathway.

**Figure 7.**
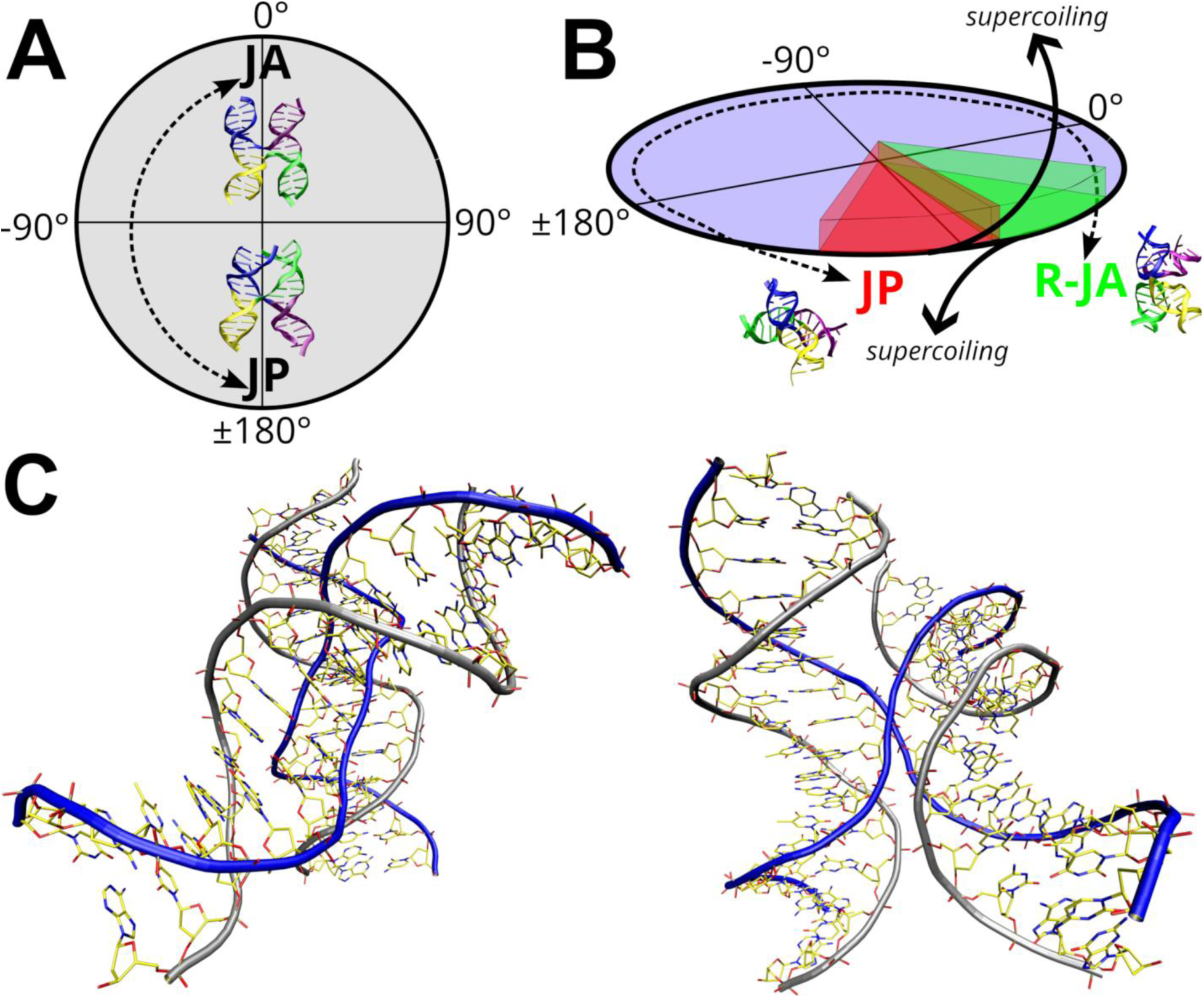
Schematic representation of the JA/JP transition of the HJ. (A) The simple model in which the JA and JP states are separated by 180° of interhelical angle rotation and related by two-fold symmetry. (B) The JA/JP transition pathway (indicated by dashed arrow line) suggested by our calculations. Rather than being diametrically opposed, the R-JA and JP states are both shifted toward the supercoiling limits (black arrows pointing above and below the circle) of the HJ DNA, resulting in a partial overlap of their interhelical angle ranges accessible during thermal fluctuations. (C) Structure of the HJ in the JP state as observed in our simulations, shown from two different viewing angles. The DNA backbone of the non-exchanging and exchanging strands is depicted as grey and blue tubes, respectively. A PDB file of the JP state of J1 obtained from the standard MD simulations is provided in the Supplementary Information.

## Conclusions

We present an *in silico* investigation of Holliday junctions (HJs), demonstrating that the parallel state (JP), which has long remained elusive in structural experiments, is structurally compatible with the HJ architecture and dynamically accessible in aqueous solution. Across extensive standard MD simulations, we observed spontaneous JA/JP transitions on the ∼10 μs timescale, demonstrating that the JP state can be reached through thermal fluctuations alone. Importantly, the helical arms remained well formed throughout the transition and within the JP state, indicating that the parallel HJ conformation is fully compatible with canonical DNA duplex geometry despite its absence from experimentally resolved structures. The transition proceeds without passage through the open HJ state and exhibits a predominantly diffusive character. Enhanced-sampling REUS calculations further characterized the thermodynamics of the JA/JP transition. In all systems examined, the experimentally observed R-JA state corresponded to the global free-energy minimum. Nevertheless, the JP state consistently formed a distinct local free-energy minimum, with the predicted free-energy difference strongly dependent on the FF employed. Using the OL21 FF, the JP state was only modestly less stable than R-JA, with a free-energy difference of ∼1–2 kcal/mol. In contrast, the more recent OL24 FF further stabilized the experimentally observed R-JA state, increasing the predicted free-energy difference to ∼4.5 kcal/mol, while preserving a well-defined JP free-energy minimum. Thus, although the exact equilibrium population of the JP state is somewhat uncertain, both FFs clearly support its existence as a kinetically stable conformational state in solution. Taken together, our results suggest that parallel Holliday junctions should be experimentally accessible in solution, even if populated only at very low levels, potentially below 1% assuming the values indicated by the OL24 FF are closer to reality. Overall, these findings highlight that FF performance is a non-trivial issue for simulations of HJs despite their relatively simple topology of four canonical B-DNA helices joined into a four-way junction. A comprehensive assessment of FF accuracy is beyond the scope of the present study and will be the subject of future work.

Several additional factors may contribute to the apparent absence of JP states from earlier experimental studies. First, the low populations of the JP state may inherently complicate direct detection. Second, JA/JP interconversion occurs on the microsecond timescale, potentially limiting its resolution by experimental techniques. Third, the dominant R-JA and JP conformations are not geometrical opposites; instead, they differ primarily in the handedness of their global supercoiling (Figure 7). As a consequence, experimental observables, such as the distances between fluorophores covalently linked to the DNA in FRET measurements, might not differ to a sufficient degree to unambiguously distinguish the R-JA and JP states.

Overall, our work provides a detailed computational perspective on the longstanding debate regarding the existence of parallel Holliday junctions in solution. While the antiparallel conformation remains unequivocally the thermodynamically favored state, our simulations identify the JP conformation as a physically robust, low-population state within the natural conformational landscape of the Holliday junction. These findings refine our understanding of HJ structural dynamics and provide a framework for future experimental efforts aimed at detecting and characterizing parallel junction conformations.

## Author Contributions

Toon Lemmens: Data curation, Formal analysis, Methodology, Investigation, Writing—original draft, Visualization. Jiří Šponer: Formal analysis, Writing—original draft, Funding acquisition. Petr Stadlbauer: Methodology, Formal analysis, Writing—original draft. Miroslav Krepl: Conceptualization, Funding acquisition, Validation, Writing—original draft, Methodology, Project administration, Supervision.

## Supplementary Data statement

Supplementary Data are available at NAR online.

## Acknowledgement

This work has been conducted in the sustainability period of the project SYMBIT No. CZ.02.1.01/0.0/0.0/15_003/0000477 as its follow-up activity. Access to CESNET storage facilities provided by the project “e-INFRA CZ” under the programme “Projects of Large Research, Development, and Innovations Infrastructures” LM2023054), is appreciated.

## Data availability

The data necessary to review and reproduce the calculations performed in this work have been deposited in Zenodo under accession code X.

