## Supplementary for "Parallel DNA Holliday Junctions: Myth or Reality? An Atomistic Molecular Dynamics Study"

### Supplementary Data

### Supplementary Tables

Table S1: **Average CV values (interhelical angle) corresponding to the R-JA, L-JA and JP states sampled in the standard MD simulations.**

| | Average CV (degrees) | | |
|---|---|---|---|
| **System[a]** | **R-JA** | **L-JA** | **JP** |
| **J1/OL21** | 36.8 (±10.7) | -56.8 (±28.8) | -254.4 (±15.0) |
| **J11-IsoI/OL21** | 34.7 (±16.0) | -41.6 (±26.6) | -250.5 (±15.6) |
| **J11-IsoII/OL21** | 37.5 (±9.4) | -59.0 (±30.6) | -253.6 (±19.7) |
| **J1-IsoI/OL24** | 42.2 (±10.7) | -34.0 (±27.6) | -243.6 (±16.9) |
| **J11-IsoI/OL24** | 25.5 (±13.1) | -28.3 (±20.1) | -244.2 (±17.3) |
| **J11-IsoII/OL24** | 43.6 (±8.9) | -73.9 (±27.9) | -244.5 (±17.5) |

[a]To calculate the averages, the data was clustered into three groups with CV values in range of [-300°, -200°], [-125°, 0°], and [0°, 100°] assigned to the JP, L-JA, and R-JA states, respectively. Note that the population distributions of these states are not converged in standard MD trajectories and the numbers stated here should therefore be interpreted with caution.

### Supplementary Figures

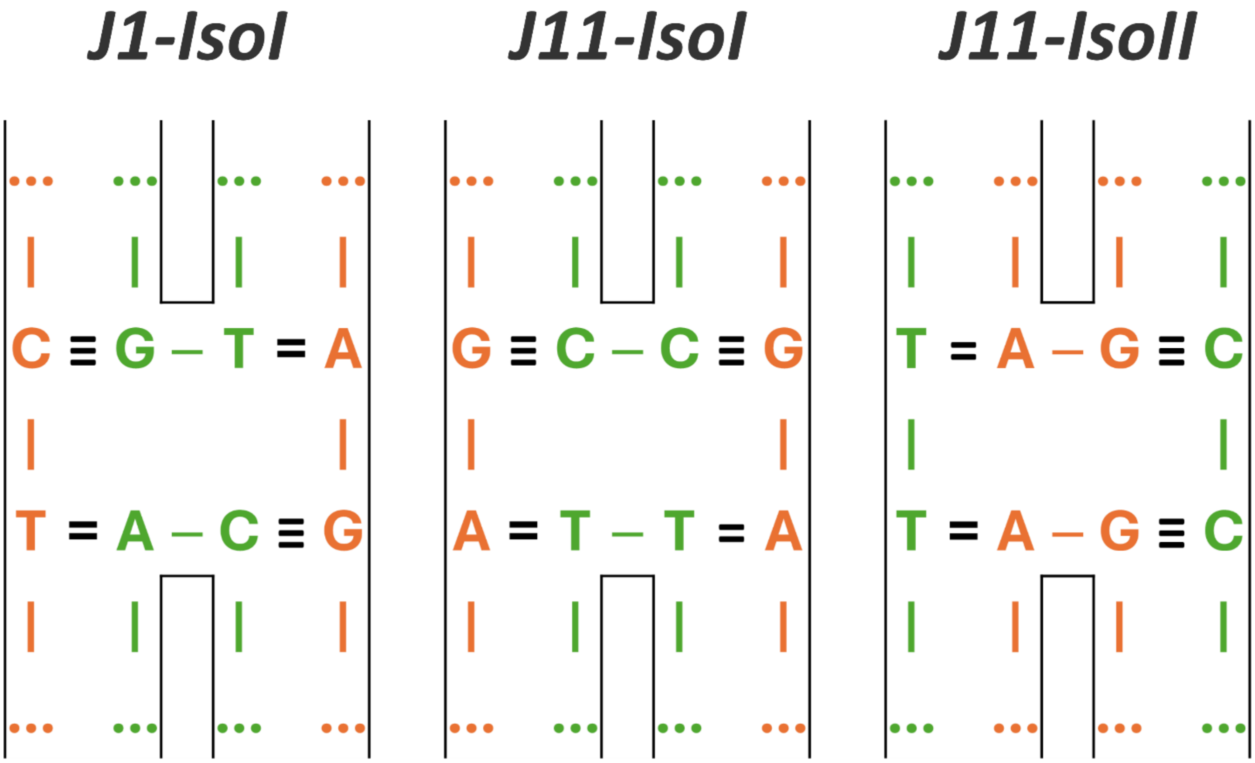


Figure S1: **Branching-point base pairs of the J1, J11-isoI, and J11-isoII junctions.** In J1 and J11-isoI, the continuous and exchanging strands are shown in orange and green, respectively. The strand assignments are reversed in J11-isoII.

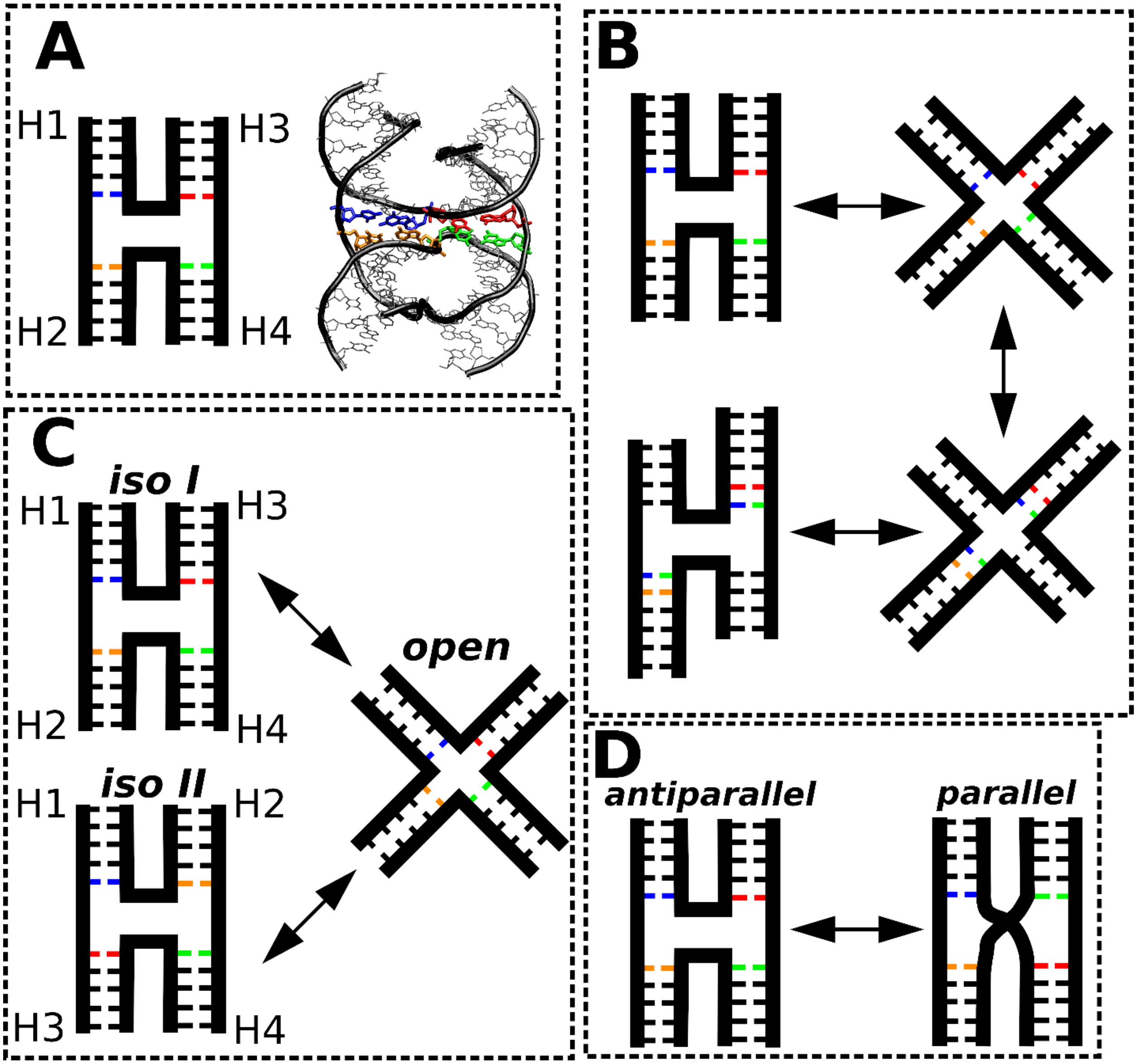


Figure S2: **Characteristic conformational transitions of a Holliday junction (HJ).** (A) Schematic representation of the HJ. Each arm is labeled with a colored base pair located in the branch region. (B) ***Branch migration***, in which the crossover point moves along the DNA duplex. The present study employed immobile HJs, thereby preventing this type of motion. (C) ***Isomerization***, involving interconversion between alternative antiparallel stacked conformations of the junction. This process requires the open state as an intermediate and was not investigated in the present study. (D) ***Antiparallel–parallel transition***, representing conversion between antiparallel and parallel HJ conformations. This transition is the focus of the present study.

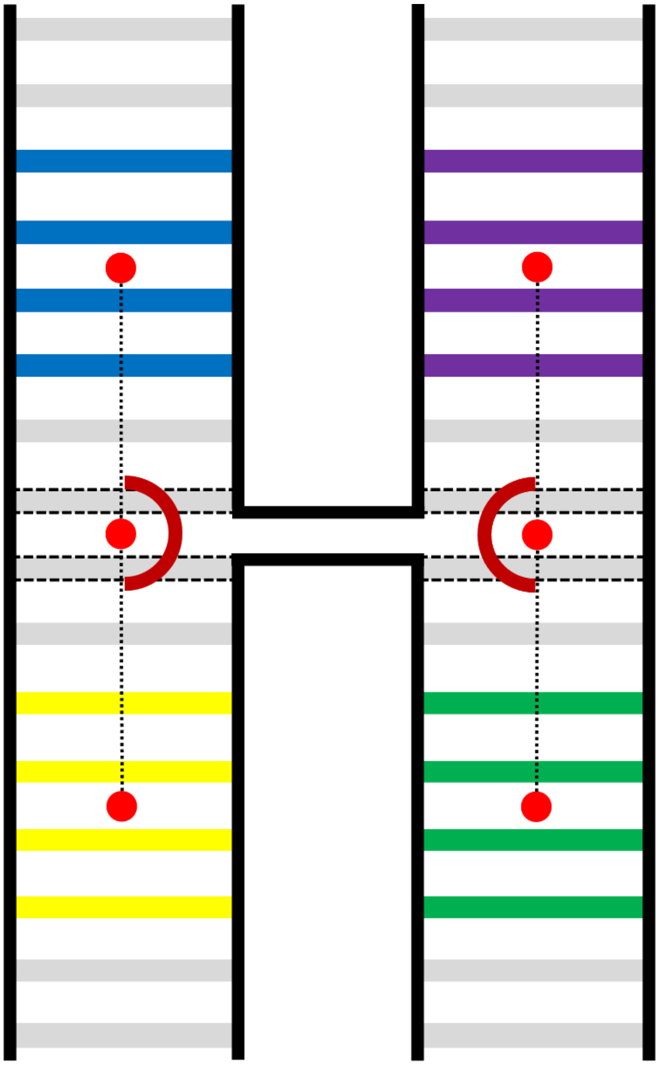

Figure S3: **Visualization of the angular restraints applied during the REUS simulations.** Each of the two restraints was defined by two centers of mass (COMs) of the C1´ atoms of the colored base pairs along a continuous HJ arm (as also defined for the CV, see main text Figure 2A). The middle points were defined with the COMs of the C1′ atoms of the four nucleotides closest to the branch point (base pairs surrounded by a dotted line). The points defining the angle are labeled with red dots and the point of rotation visualized with dark-red semi-circle. These restraints weren't applied in any of the standard MD simulations.

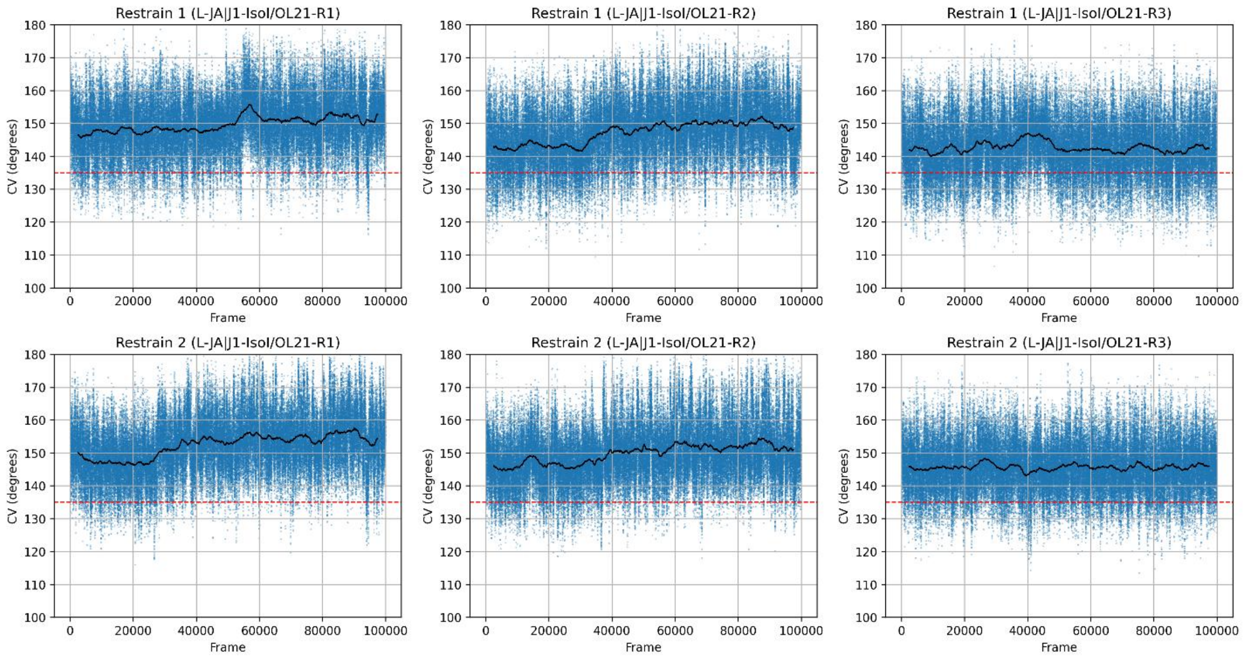


Figure S4: **Bending of the continuous helices in selected standard MD simulations.** An example is shown for the J1/OL21 system initiated from state L-JA. For definition of the bending see main text Methods and Figure S3. Black line represents the running average of the angle value while the red dotted line represents the lower-wall restraint applied in the REUS simulations.

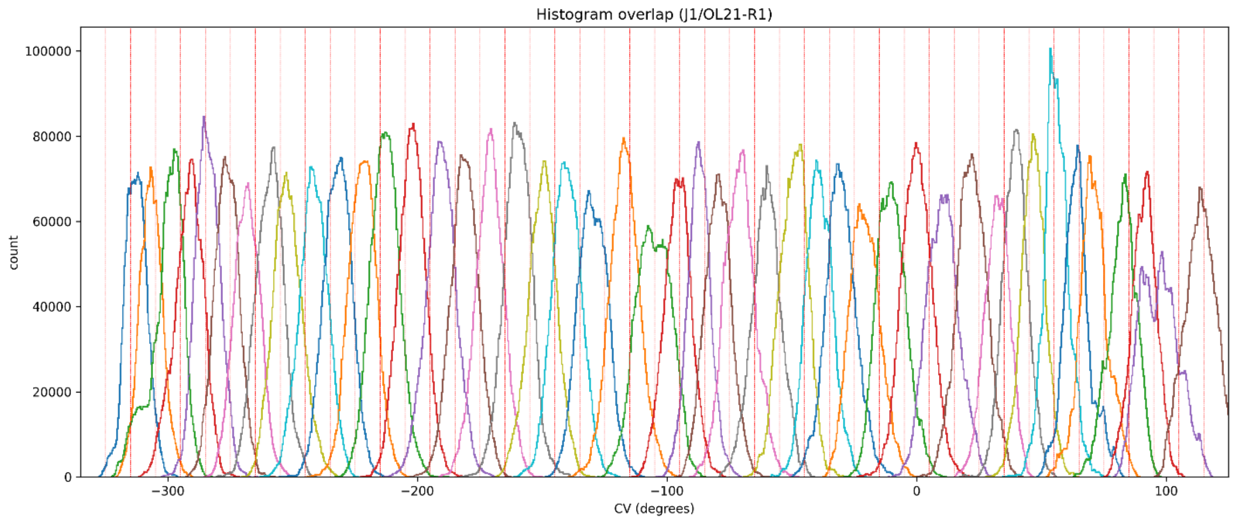
Histogram overlap (J1/OL21-R1)
count
0
20000
40000
60000
80000
100000
−300
−200
−100
0
100
CV (degrees)

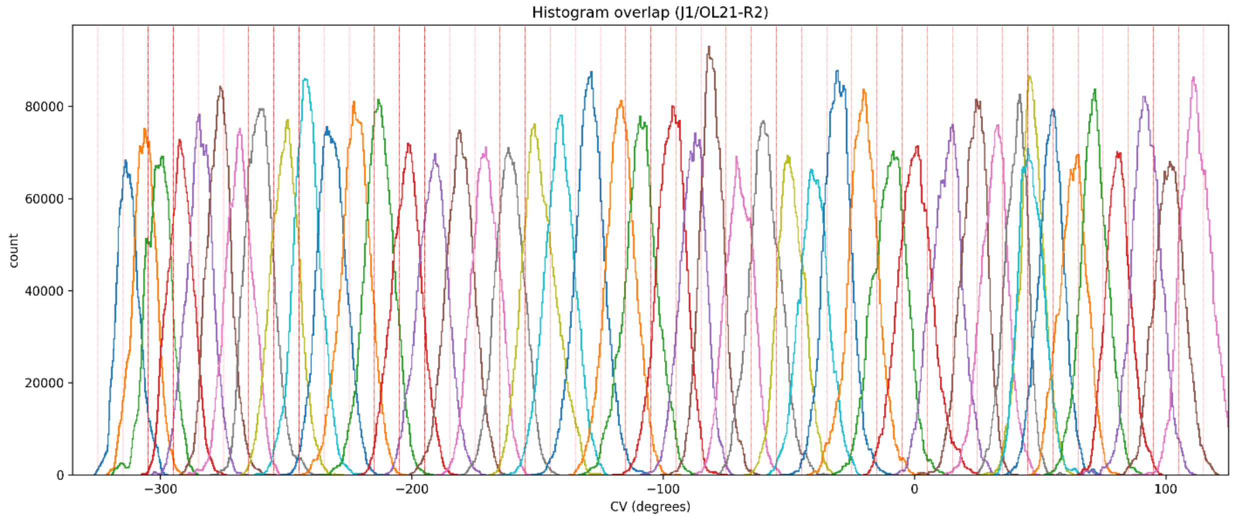
Histogram overlap (J1/OL21-R2)
count
0
20000
40000
60000
80000
−300
−200
−100
0
100
CV (degrees)

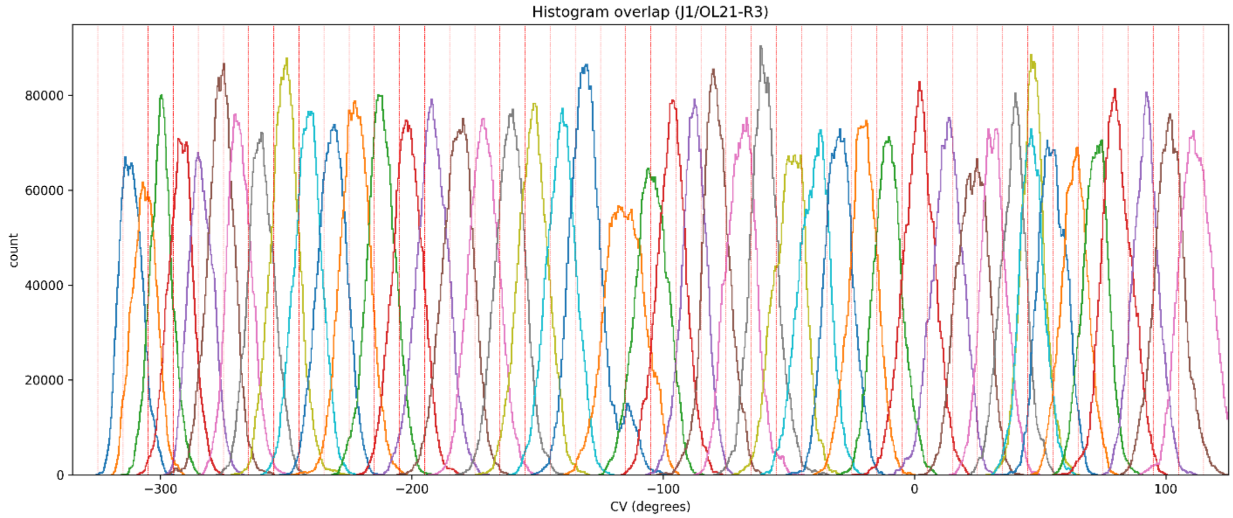
Histogram overlap (J1/OL21-R3)
count
0
20000
40000
60000
80000
−300
−200
−100
0
100
CV (degrees)

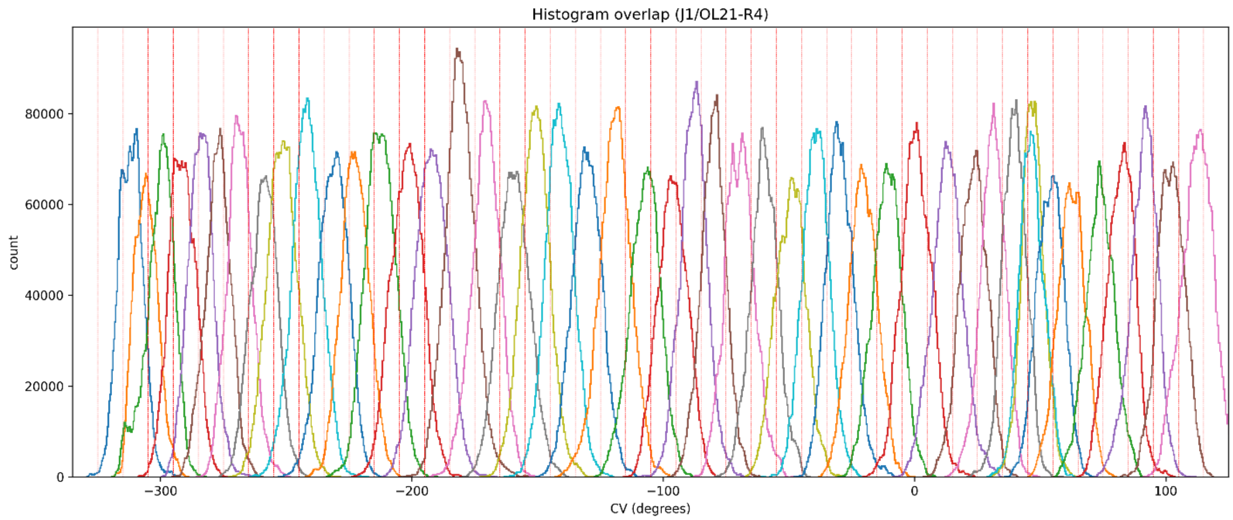


Figure S5: **Histograms of the CV value in REUS windows in the basic replica of the J1/OL21 system.** The vertical red lines define the boundaries of the window with the target value in the center. All calculated replicates are shown.

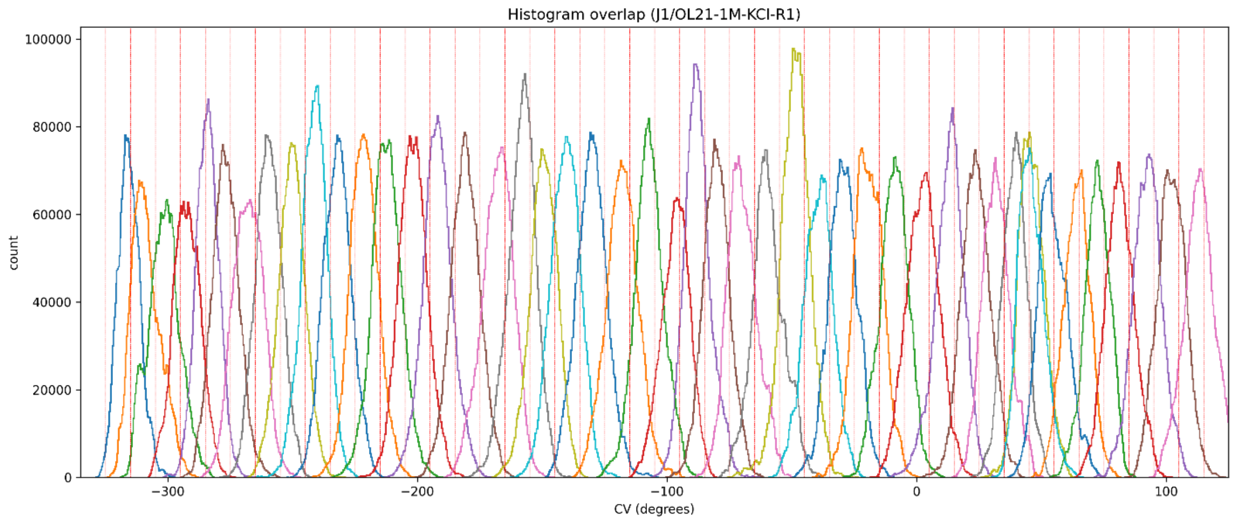
Histogram overlap (J1/OL21-1M-KCl-R1)
count
CV (degrees)
0
20000
40000
60000
80000
100000
−300
−200
−100
0
100

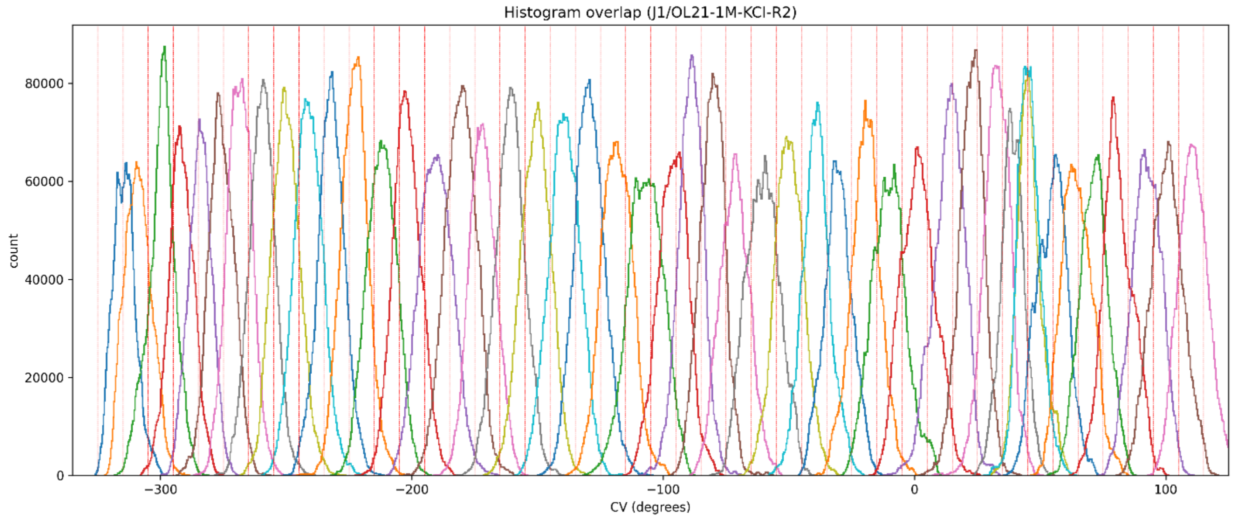
Histogram overlap (J1/OL21-1M-KCl-R2)
count
CV (degrees)
0
20000
40000
60000
80000
−300
−200
−100
0
100

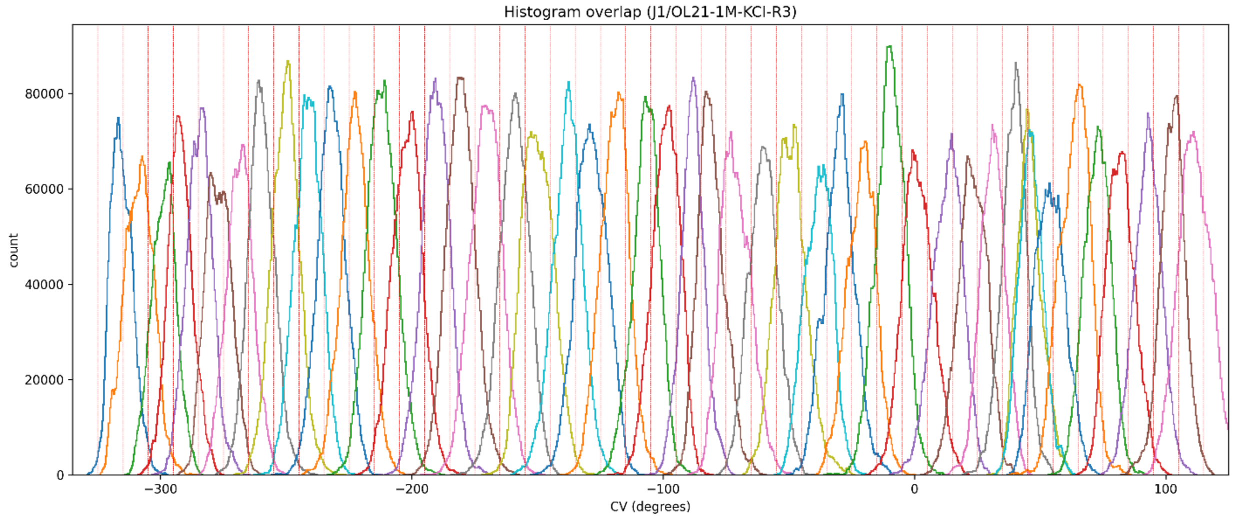
Histogram overlap (J1/OL21-1M-KCl-R3)
count
CV (degrees)
0
20000
40000
60000
80000
−300
−200
−100
0
100

Figure S6: **Histograms of the CV value in REUS windows in the basic replica of the J1/OL21-1M-KCl system.** The vertical red lines define the boundaries of the window with the target value in the center. All calculated replicates are shown.

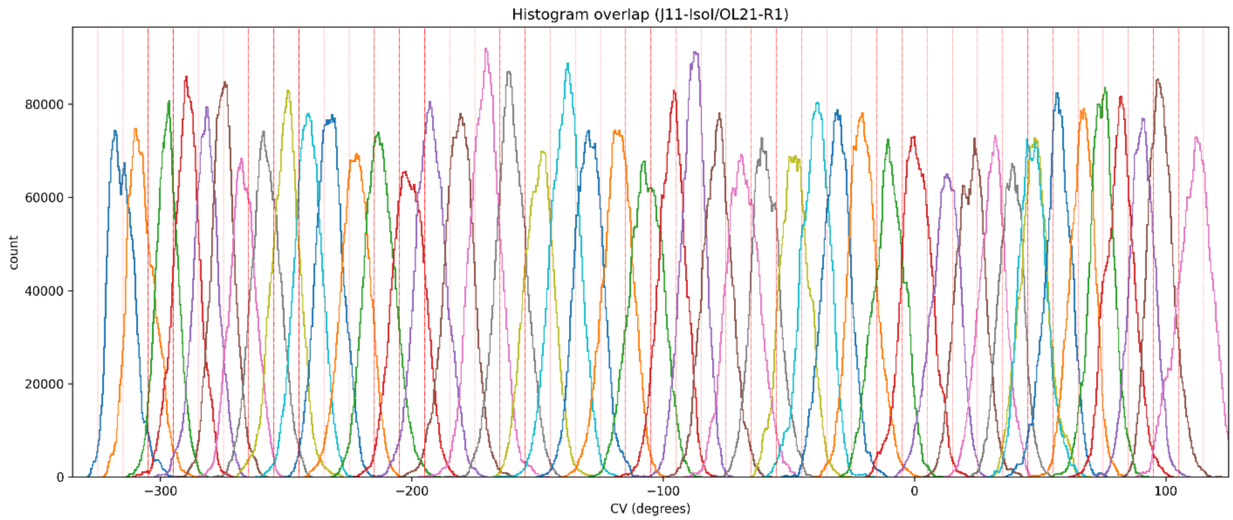
Histogram overlap (J11-IsoI/OL21-R1)
count
CV (degrees)

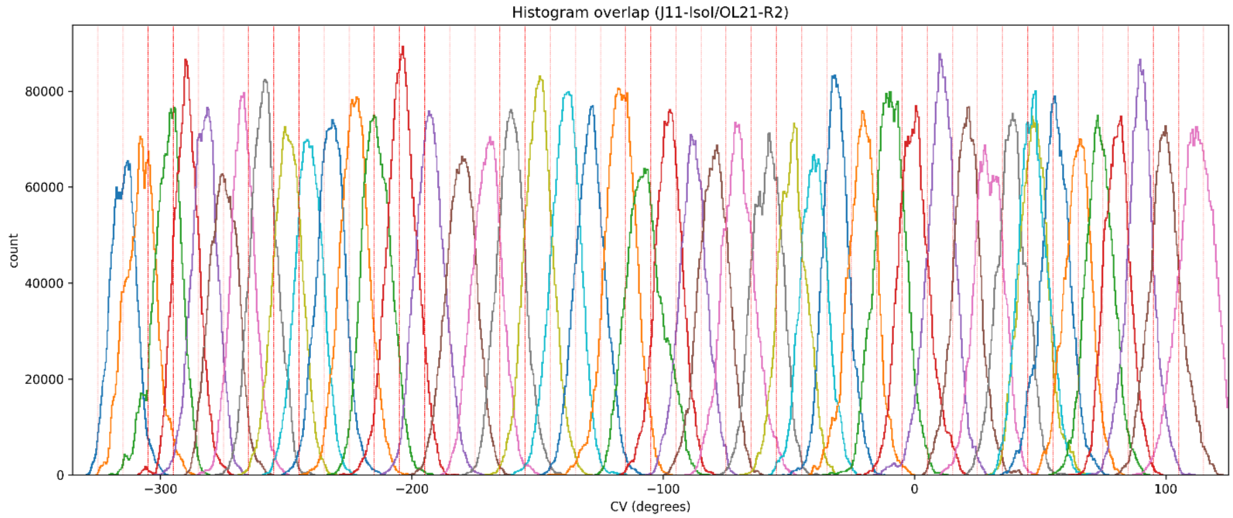
Histogram overlap (J11-IsoI/OL21-R2)
count
CV (degrees)

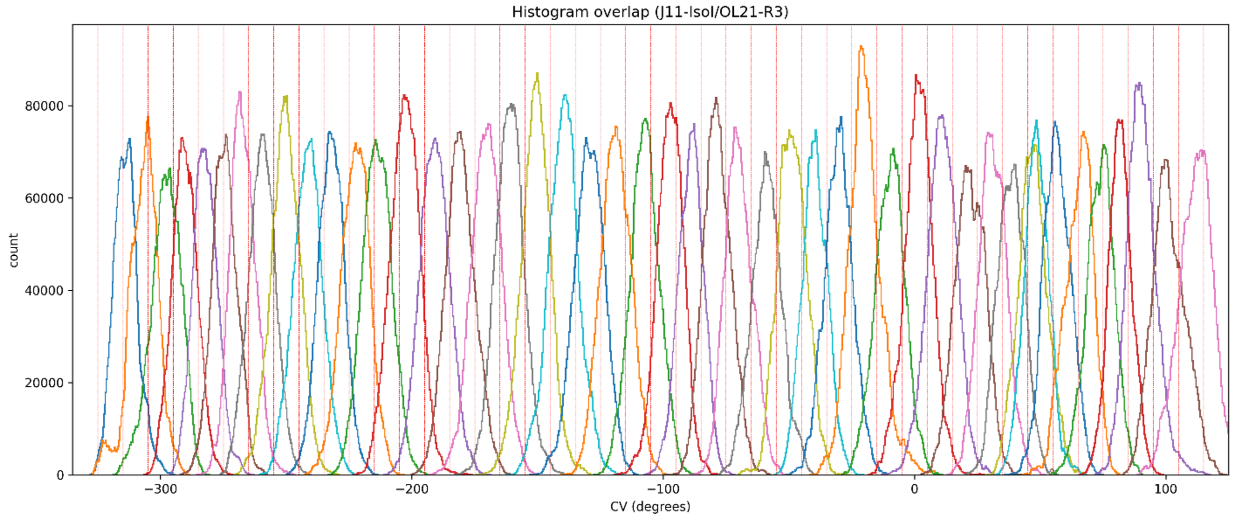
Histogram overlap (J11-IsoI/OL21-R3)
count
CV (degrees)

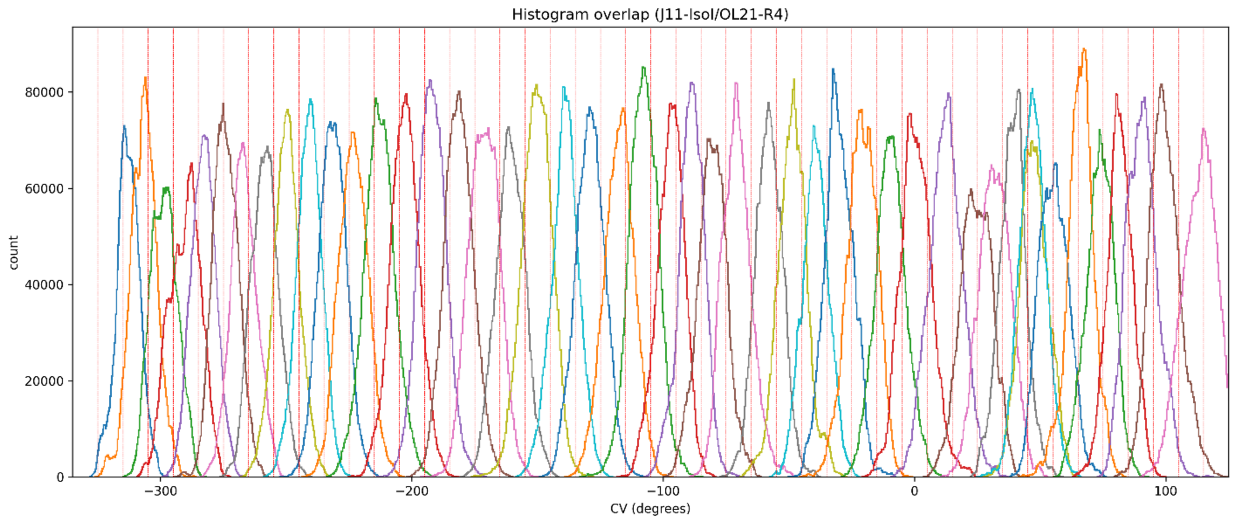


Figure S7: **Histograms of the CV value in REUS windows in the basic replica of the J11-IsoI/OL21 system.** The vertical red lines define the boundaries of the window with the target value in the center. All calculated replicates are shown.

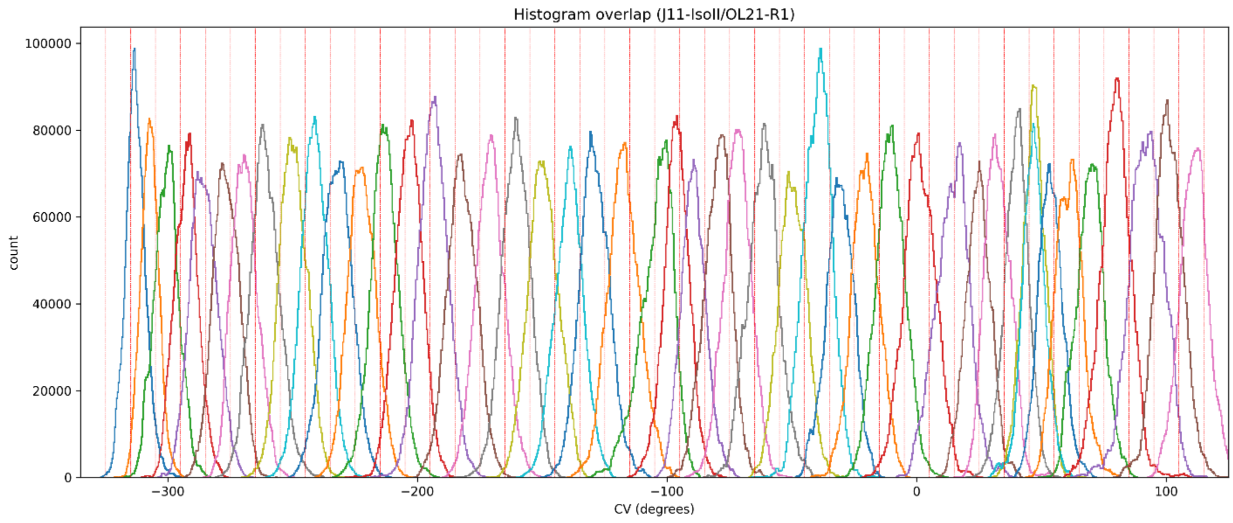
Histogram overlap (J11-IsoII/OL21-R1)
count
CV (degrees)

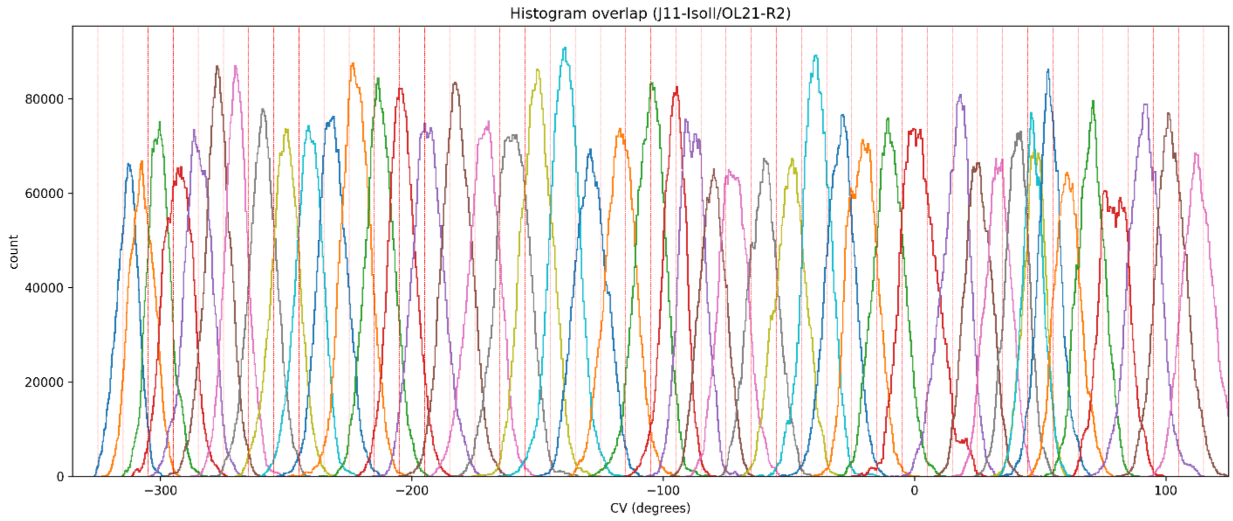
Histogram overlap (J11-IsoII/OL21-R2)
count
CV (degrees)

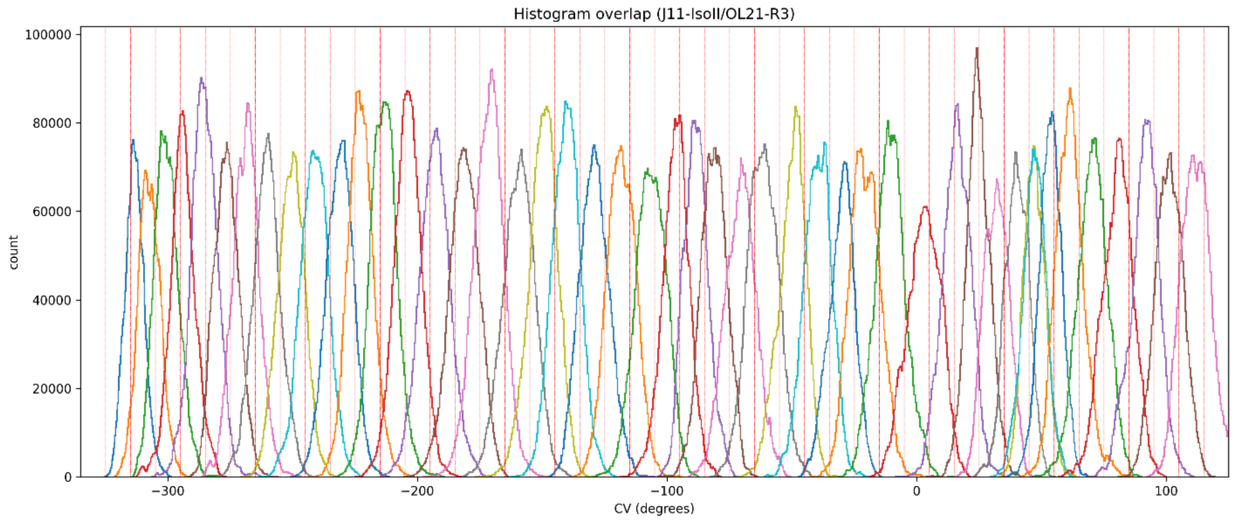
Histogram overlap (J11-IsoII/OL21-R3)
count
CV (degrees)

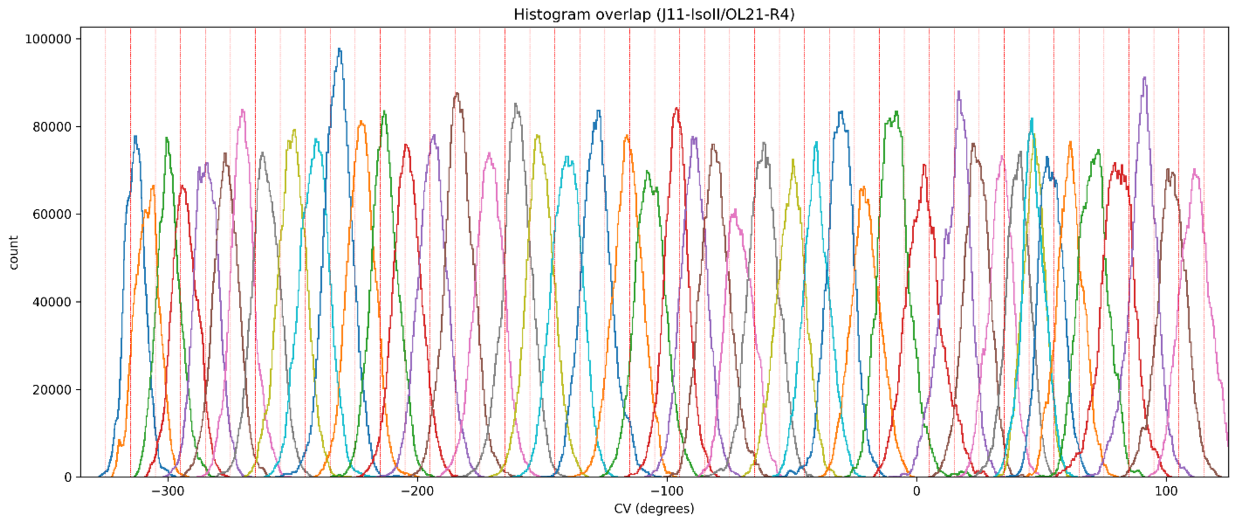


Figure S8: **Histograms of the CV value in REUS windows in the basic replica of the J11-IsoII/OL21 system.** The vertical red lines define the boundaries of the window with the target value in the center. All calculated replicates are shown.

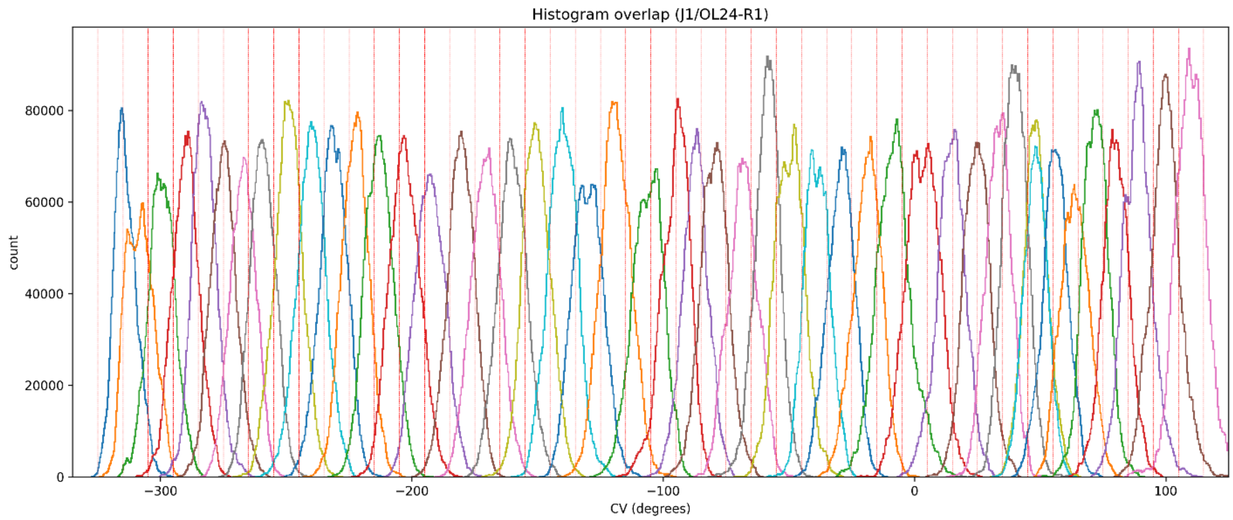
Histogram overlap (J1/OL24-R1)
count
CV (degrees)

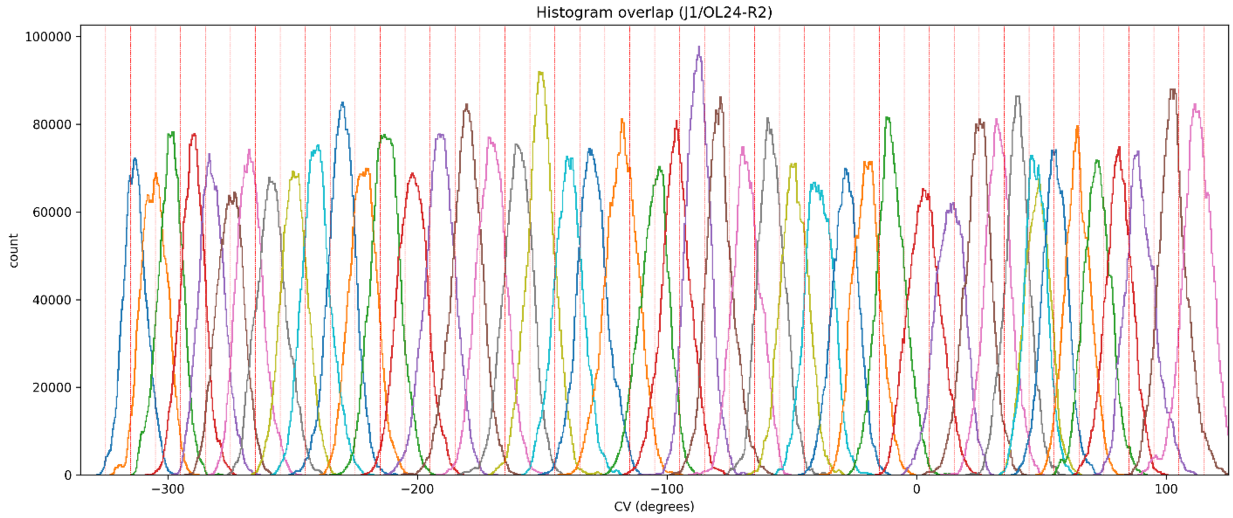
Histogram overlap (J1/OL24-R2)
count
CV (degrees)

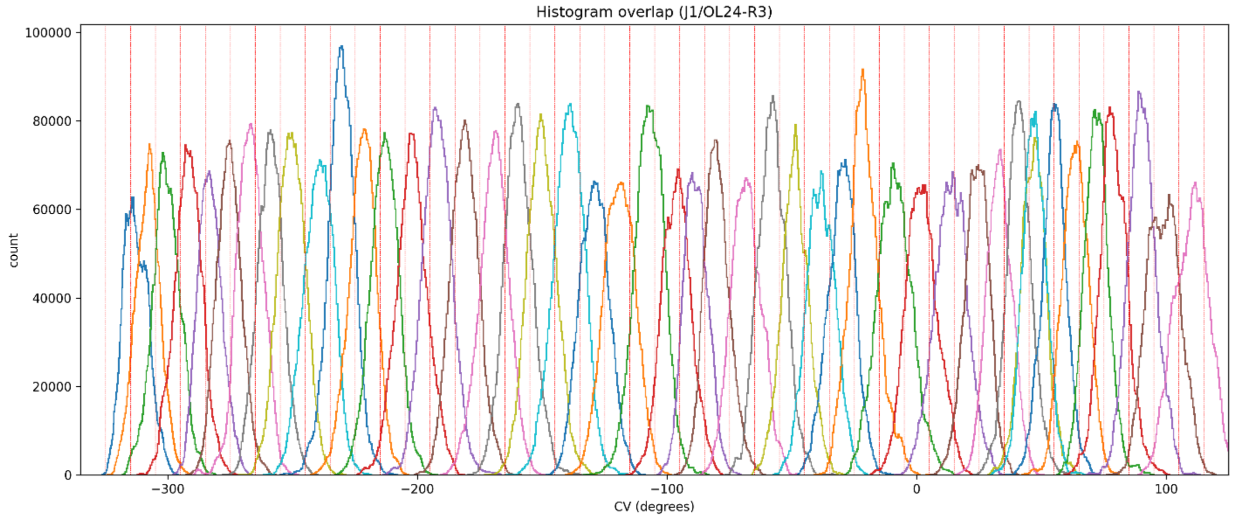
Histogram overlap (J1/OL24-R3)
count
CV (degrees)

Figure S9: **Histograms of the CV value in REUS windows in the basic replica of the J1/OL24 system.** The vertical red lines define the boundaries of the window with the target value in the center. All calculated replicates are shown.

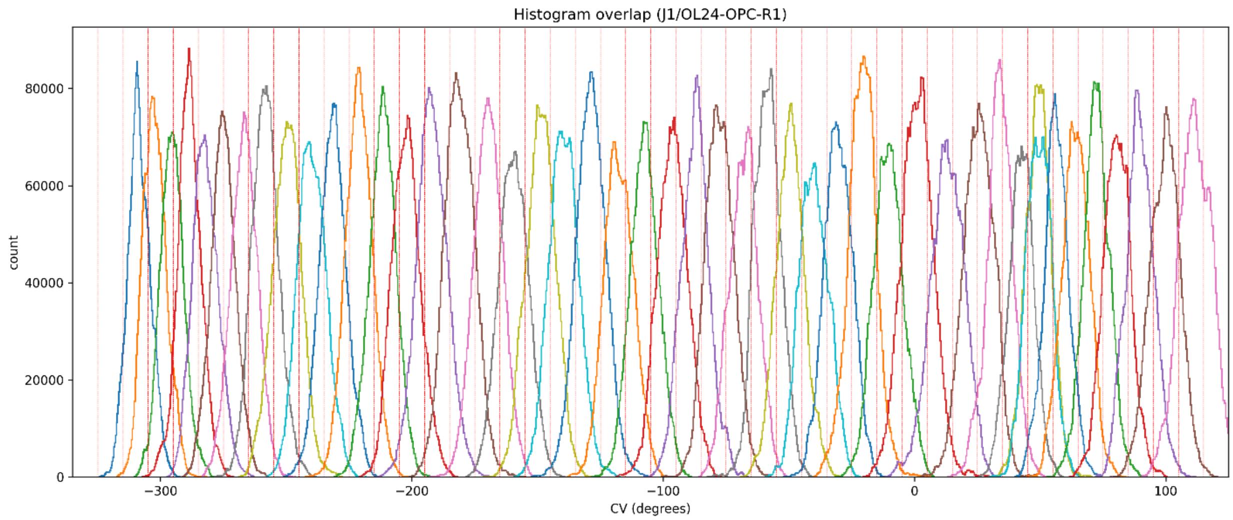

Histogram overlap (J1/OL24-OPC-R1)
count
CV (degrees)


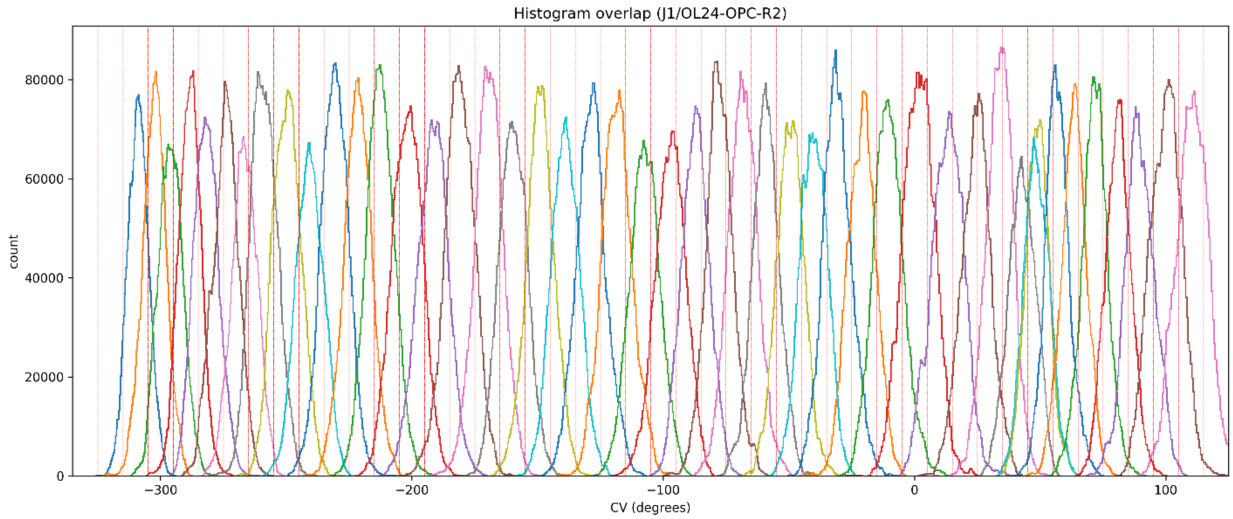

Histogram overlap (J1/OL24-OPC-R2)
count
CV (degrees)


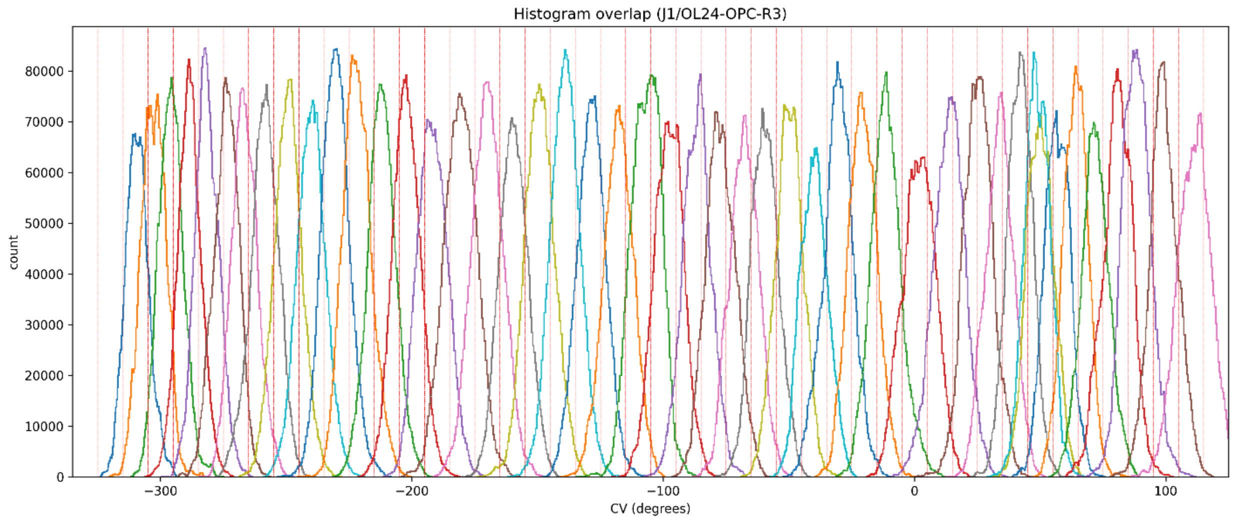

Histogram overlap (J1/OL24-OPC-R3)
count
CV (degrees)

Figure S10: **Histograms of the CV value in REUS windows in the basic replica of the J1/OL24-OPC system.** The vertical red lines define the boundaries of the window with the target value in the center. All calculated replicates are shown.

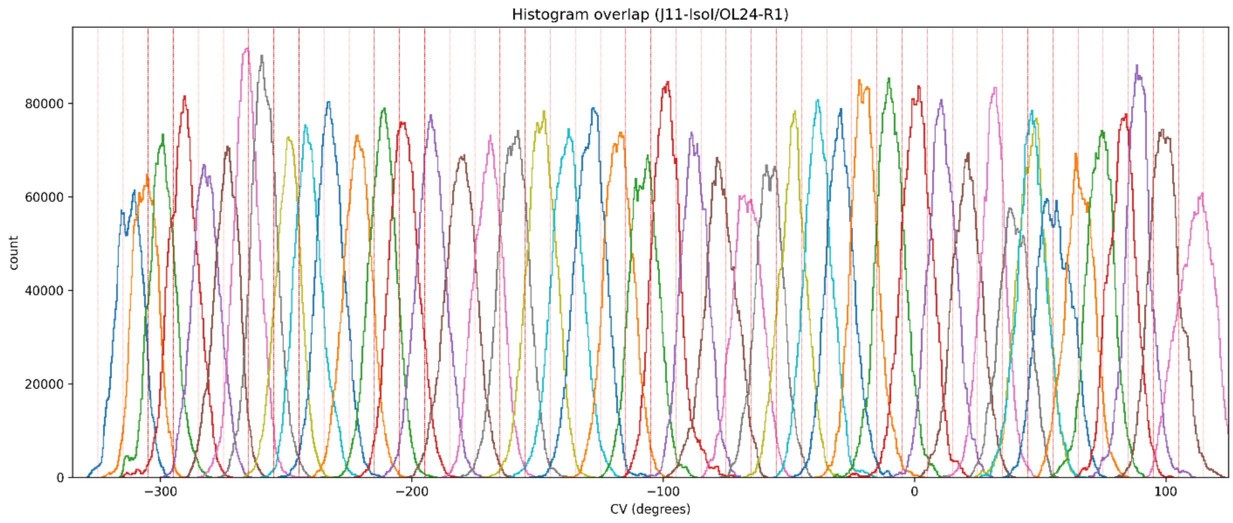
Histogram overlap (J11-IsoI/OL24-R1)
count
CV (degrees)

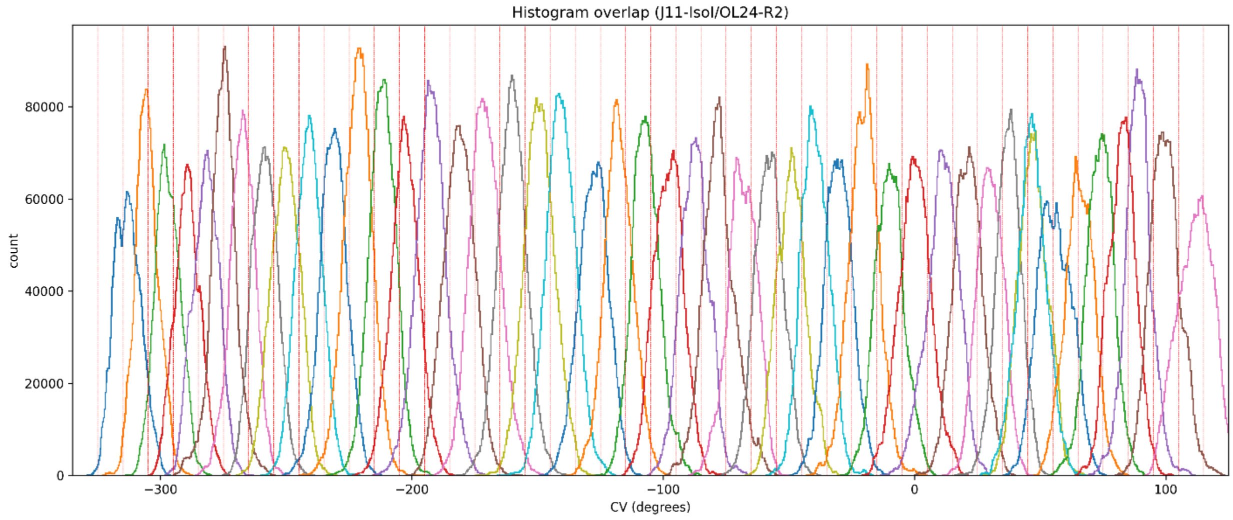
Histogram overlap (J11-IsoI/OL24-R2)
count
CV (degrees)

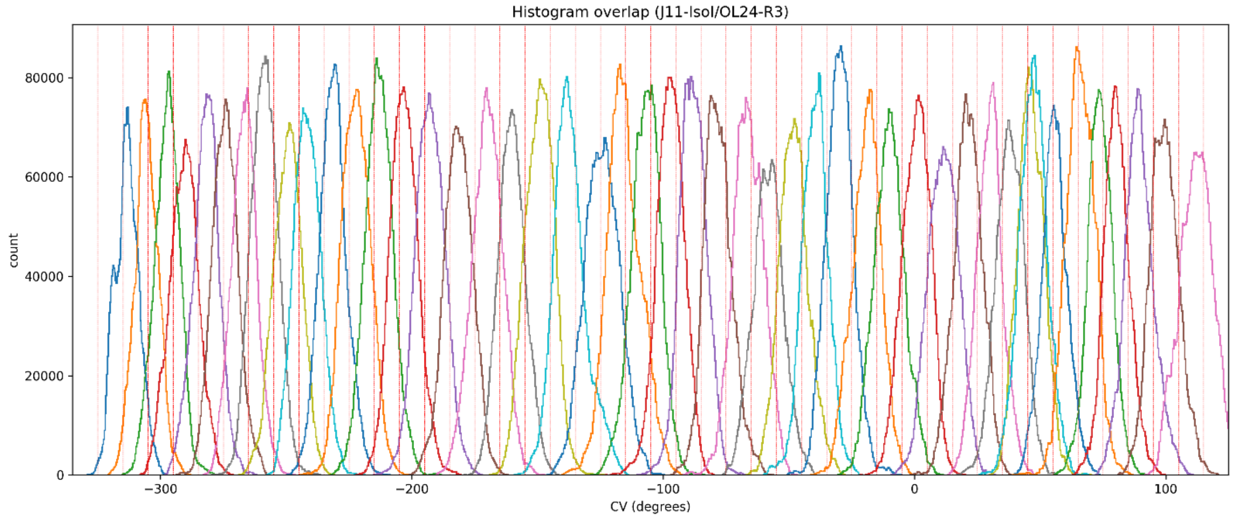
Histogram overlap (J11-IsoI/OL24-R3)
count
CV (degrees)

Figure S11: **Histograms of the CV value in REUS windows in the basic replica of the J11-IsoI/OL24 system.** The vertical red lines define the boundaries of the window with the target value in the center. All calculated replicates are shown

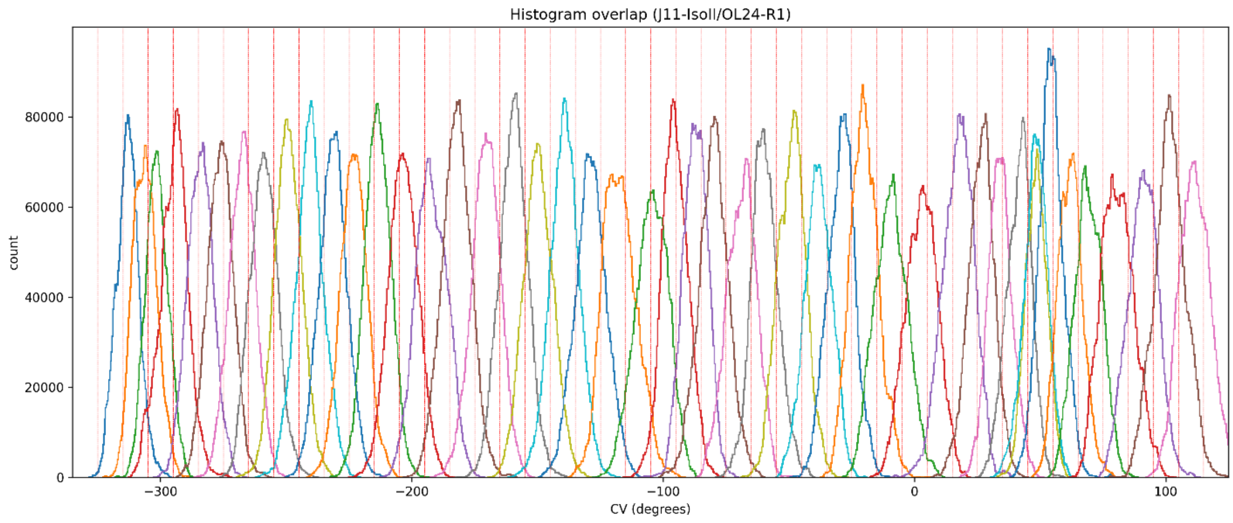
Histogram overlap (J11-IsoII/OL24-R1)
count
CV (degrees)

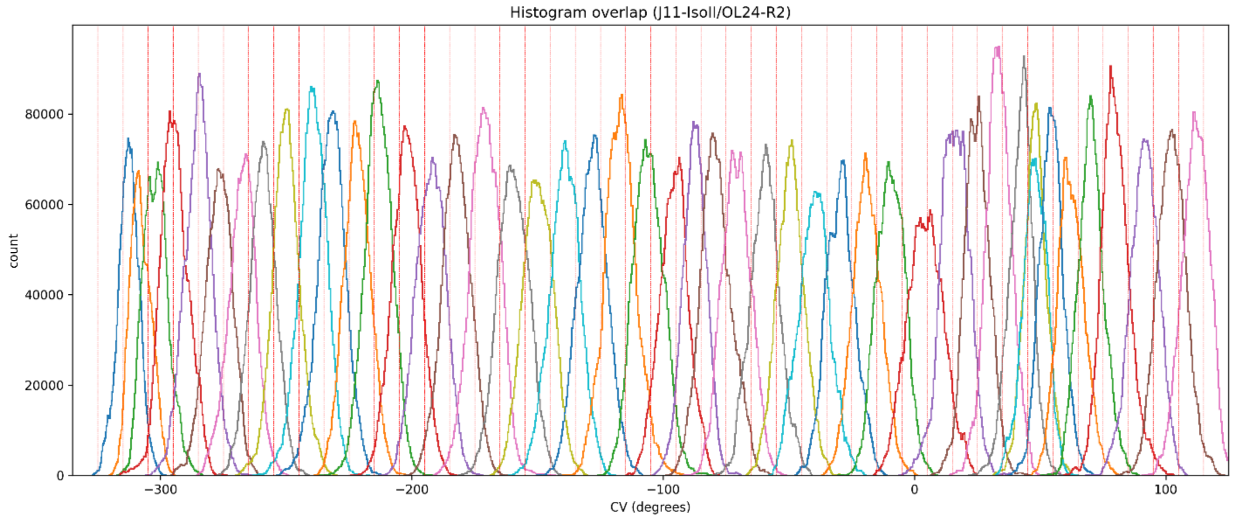
Histogram overlap (J11-IsoII/OL24-R2)
count
CV (degrees)

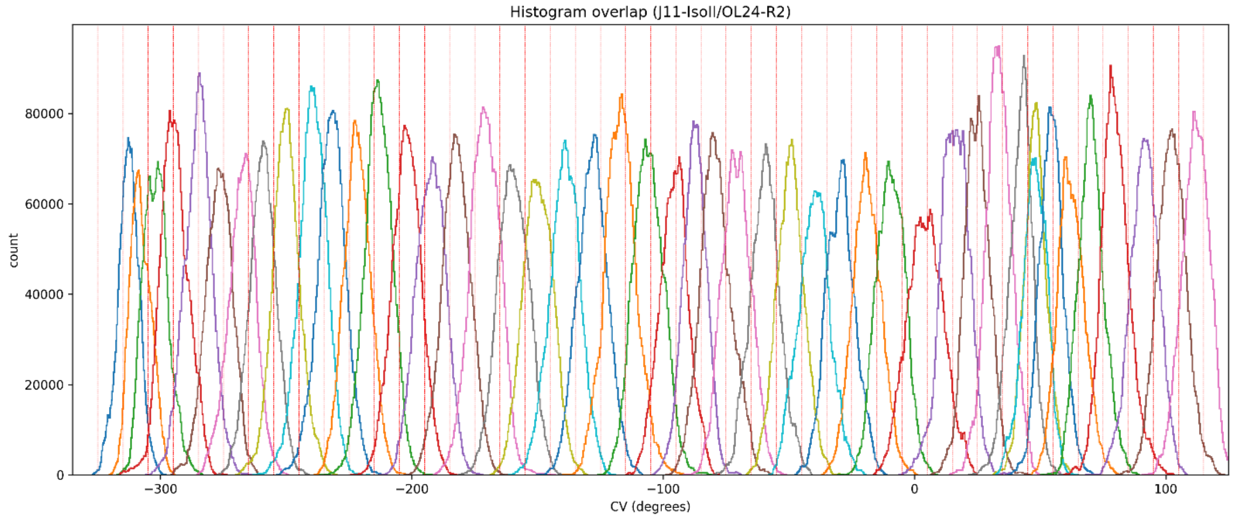
Histogram overlap (J11-IsoII/OL24-R2)
count
CV (degrees)

Figure S12: **Histograms of the CV value in REUS windows in the basic replica of the J11-IsoII/OL24 system.** The vertical red lines define the boundaries of the window with the target value in the center. All calculated replicates are shown

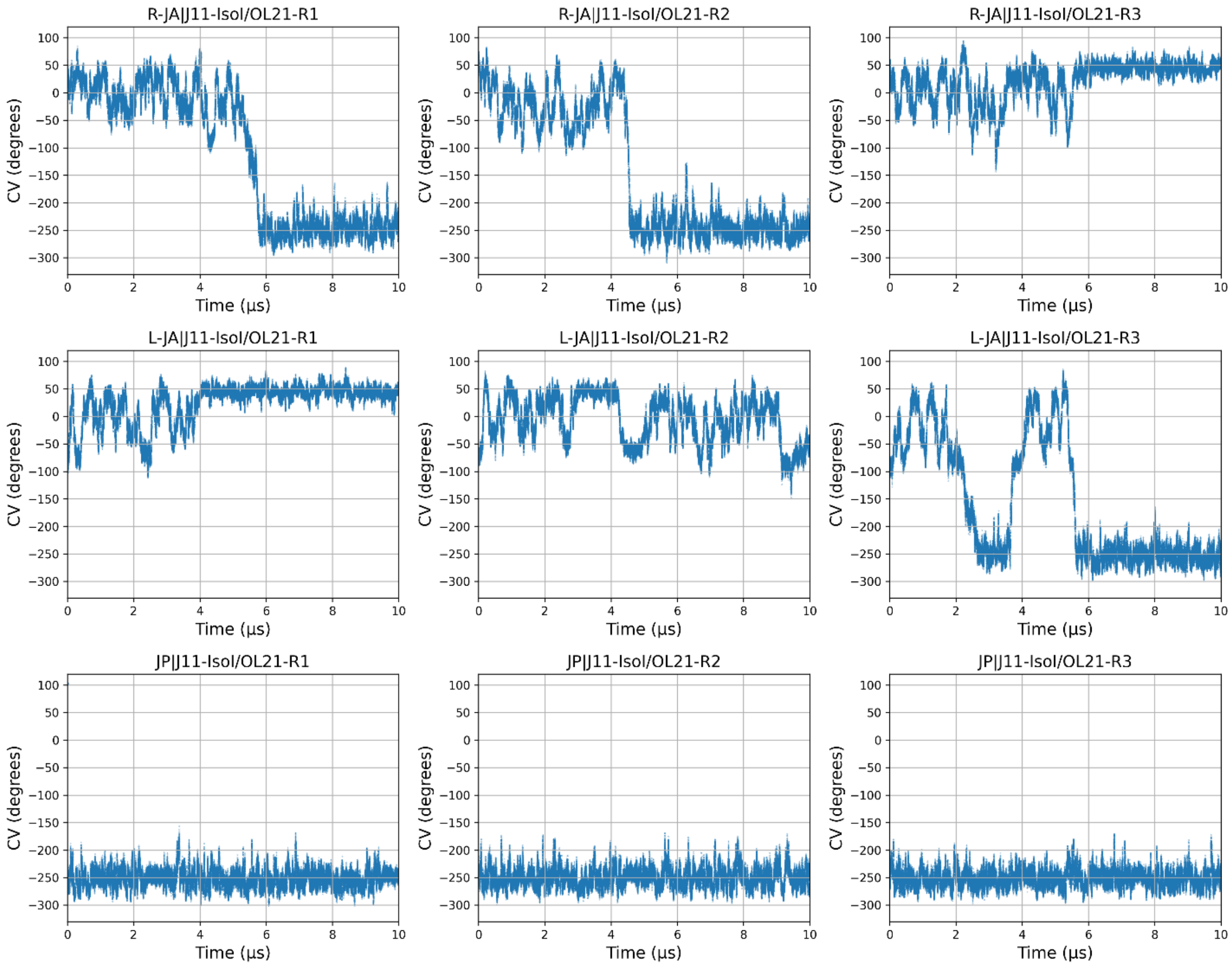


Figure S13: **Time evolution of the CV values sampled in standard MD simulations of the J11-IsoI/OL21 system.** Top, middle and bottom rows show the three standard MD simulation replicates initialized from the R-JA, L-JA and JP states, respectively.

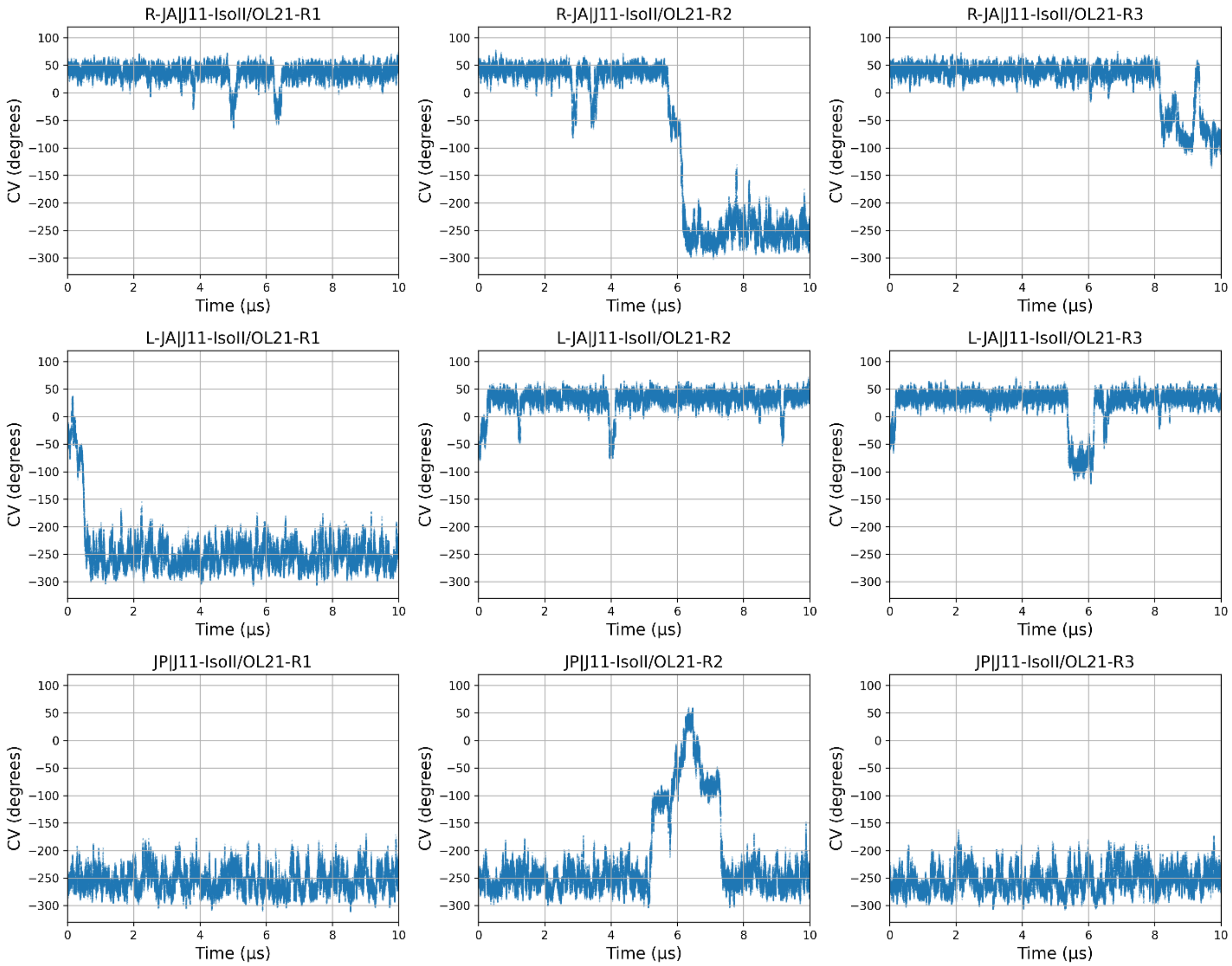


Figure S14: **Time evolution of the CV values sampled in standard MD simulations of the J11-IsoII/OL21 system.** Top, middle and bottom rows show the three standard MD simulation replicates initialized from the R-JA, L-JA and JP states, respectively.

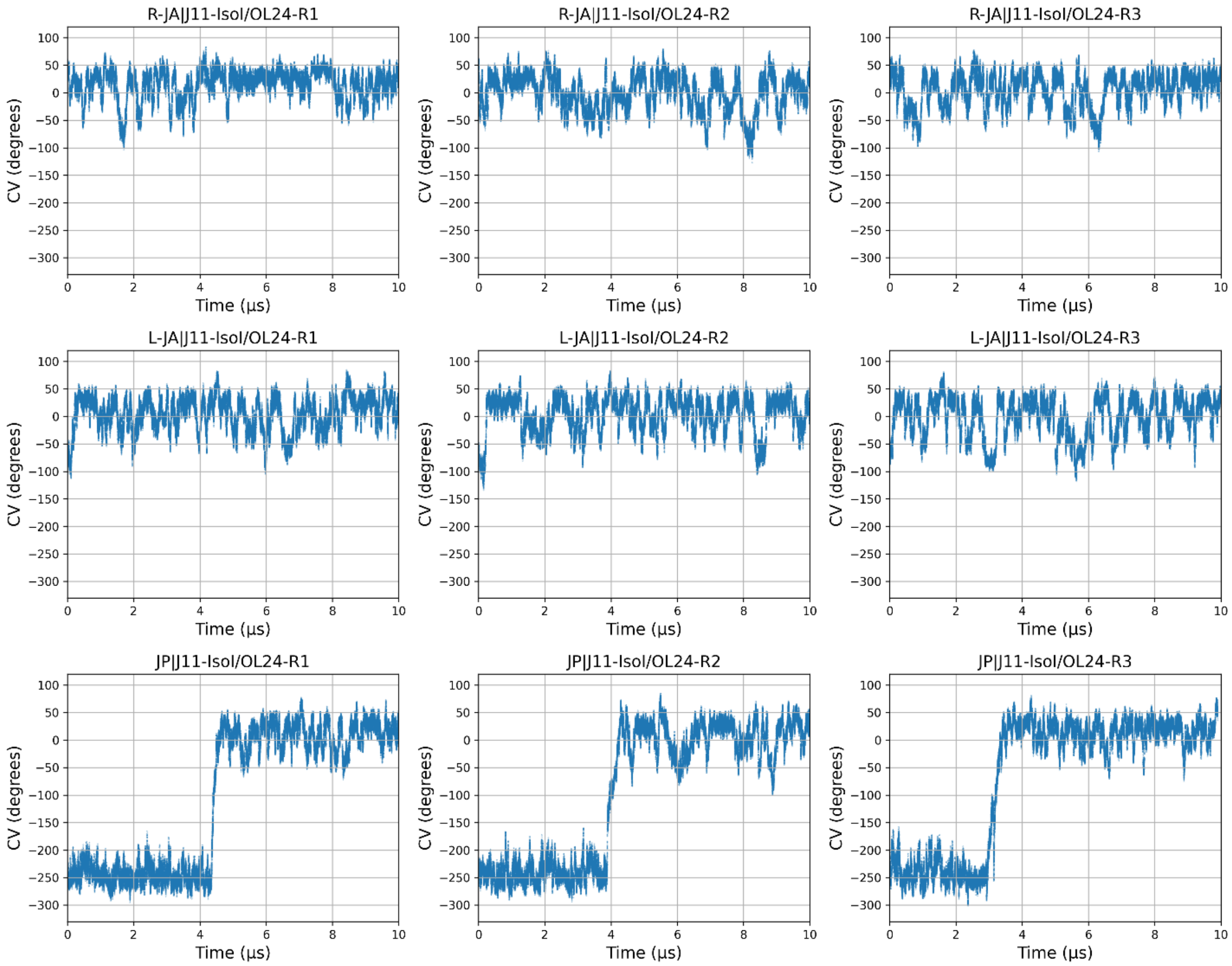


Figure S15: **Time evolution of the CV values sampled in standard MD simulations of the J11-IsoI/OL24 system.** Top, middle and bottom rows show the three standard MD simulation replicates initialized from the R-JA, L-JA and JP states, respectively.

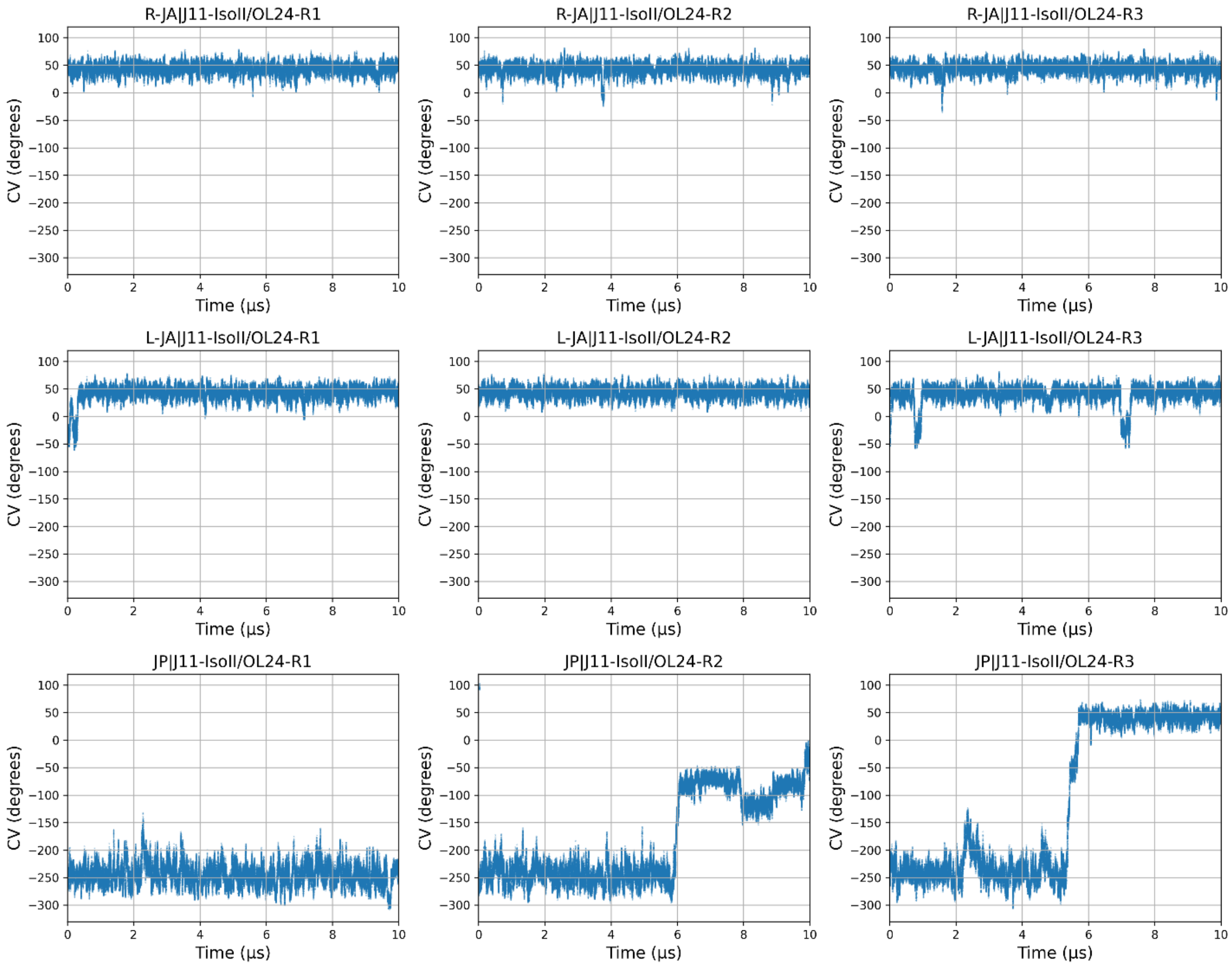


Figure S16: **Time evolution of the CV values sampled in standard MD simulations of the J11-IsoII/OL24 system.** Top, middle and bottom rows show the three standard MD simulation replicates initialized from the R-JA, L-JA and JP states, respectively.

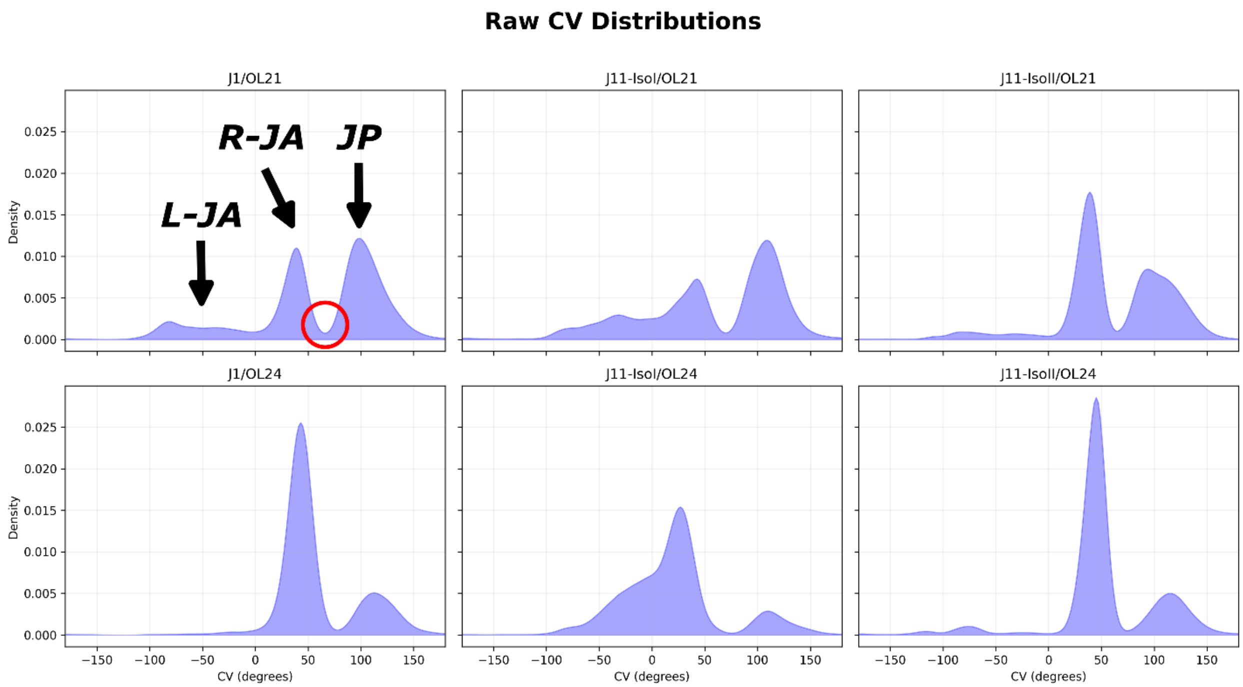


Figure S17: **Distribution of the CV values sampled in standard MD simulations without unwrapping the coordinate corresponding to the interhelical angle.** Peaks corresponding to the L-JA, R-JA, and JP states are indicated by arrows in the first graph. When the dihedral angle is not unwrapped to account for HJ supercoiling, the R-JA and JP peaks are located close to each other and exhibit a small but non-negligible overlap (indicated with a red circle).
